# Cortical responses to vagus nerve stimulation are modulated by brain state in non-human primates

**DOI:** 10.1101/2021.01.11.426280

**Authors:** Irene Rembado, Weiguo Song, David K. Su, Ariel Levari, Larry E. Shupe, Steve Perlmutter, Eberhard Fetz, Stavros Zanos

## Abstract

Vagus nerve stimulation (VNS) has been tested as therapy for several brain disorders and as a means to modulate cortical excitability and brain plasticity. Cortical effects of VNS, manifesting as vagal-evoked potentials (VEPs), are thought to arise from activation of ascending cholinergic and noradrenergic systems. However, it is unknown whether those effects are modulated by brain state at the time of stimulation. In 2 freely behaving macaque monkeys, we delivered trains of left cervical VNS at different frequencies (5-300 Hz) while recording local field potentials (LFPs) from sites in contralateral prefrontal, sensorimotor and parietal cortical areas. Brain states were inferred from spectral components of LFPs and the presence of overt movement: active awake, resting awake, REM sleep and NREM sleep. VNS elicited VEPs comprising early (<70 ms), intermediate (70-250 ms) and late (>250 ms) components in all sampled cortical areas. The magnitude of only the intermediate and late components was modulated by brain state and pulsing frequency. These findings have implications for the role of ongoing cortical activity and brain state in shaping cortical responses to peripheral stimuli, for the modulation of vagal interoceptive signaling by cortical states, and for the calibration of VNS therapies.

## Introduction

Vagus nerve stimulation (VNS) is a non-pharmacological, FDA-approved treatment for epilepsy and depression and has been tested as a possible therapy for tinnitus (Tyler R et al. 2017), post-traumatic stress disorder (Noble LJ et al. 2017), headache, sleep disorders and neurorehabilitation after stroke (Elger G et al. 2000; George MS et al. 2002; Henry TR 2002; Dawson J et al. 2016). VNS has beneficial effects on cognition and behavior, as it enhances cognitive abilities in Alzheimer’s patients (Sjogren MJ et al. 2002), and facilitates decision-making in animals (Cao B et al. 2016). The afferent vagus projects directly or indirectly to several brainstem and forebrain nuclei including the nucleus of solitary tract, the locus coeruleus and the nucleus basalis (Henry TR 2002; Hassert DL et al. 2004; Cheyuo C et al. 2011) and from there to numerous subcortical and cortical areas (Pritchard TC et al. 2000; Henry TR 2002; Collins L et al. 2021), releasing mostly norepinerphrine and acetylocholine. VNS suppresses cortical excitability (Zagon A and AA Kemeny 2000; Di Lazzaro V et al. 2004; Nichols JA et al. 2011) and enhances plasticity by facilitating reorganization of cortical maps (Porter BA et al. 2012; Shetake JA et al. 2012; Engineer CT et al. 2015). Moreover, due to the precise control of the timing of its delivery, VNS is a candidate for delivering temporary precise, closed-loop neuromodulation to the brain (Zanos S 2019), to treat neurological disorders and augment learning (Engineer ND et al. 2011; Hays SA et al. 2016; Pruitt DT et al. 2016).

Afferent volleys elicited by VNS give rise to vagal evoked potentials (VEPs) at different levels, including brainstem, hippocampus, thalamus and cerebral cortex (Car A et al. 1975; Hammond EJ et al. 1992, 1992; Ito S and AD Craig 2005). Cortical VEPs in humans are widespread and bilateral and comprise a component within the first 10-20 ms post-stimulus (Hammond EJ *et al*. 1992) and additional components with longer latencies (Upton AR et al. 1991). There are well-documented differences, with regards to features of VEPs, between healthy subjects of different ages (Fallgatter AJ et al. 2005), between healthy subjects and patients (Polak T et al. 2007), and within populations of patients with cognitive impairment (Metzger FG et al. 2012). Cortical responses to VNS may be used to screen subjects for neurodegenerative diseases, to assist the diagnosis and to track disease progression (Metzger FG *et al*. 2012; Polak T et al. 2017). Furthermore, EEG markers, like VEPs, could be used to predict effectiveness of VNS therapy (Ravan M et al. 2017).

Such uses for VEPs are confounded by factors like stimulus intensity, pulse width and pulsing frequency (Polak T et al. 2009; Hagen K et al. 2014). In addition, since VEPs are elicited by subcortical projections to the cortex and generated by stimulus-evoked cortical activity, ongoing brain activity, both cortical and subcortical, at the time of stimulation, may affect VEPs. Brain-wide oscillatory cortical activity changes between behavioral states, in both natural conditions (e.g. wakefulness and sleep cycles) and in pathological conditions (Steriade M et al. 1993; Steriade M 1997; Destexhe A et al. 1999; Vesuna S et al. 2020). Profound changes in cortical responsiveness to sensory stimuli have been documented in association with different patterns of ongoing cortical activity in awake and sleep states (Livingstone MS and DH Hubel 1981; Massimini M et al. 2005; Hennevin E et al. 2007). Likewise, cortical potentials evoked by electrical stimulation of somatic peripheral nerves have been shown to be modulated by brain states (Shaw FZ et al. 2006). However, it is unknown whether VEPs are also modulated by brain states. Given the widespread afferent vagal projections to many subcortical and cortical sites, such modulation could offer insights into the central processing and integration of “interoceptive” signals conveyed to the brain by the vagus (Paciorek A and L Skora 2020). In turn, it could have implications for how vagal interoceptive signals are modulated by ongoing brain activity in processes like emotion (Critchley HD and SN Garfinkel 2017), cognition (Tsakiris M and H Critchley 2016), decision making (Seth AK 2013) and action (Marshall AC et al. 2018), and in mental diseases in which interoception has been implicated (Khalsa SS et al. 2018).

To investigate whether and how VEPs are modulated by brain states inferred by ongoing cortical activity, we used an autonomous portable computer, the Neurochip (Zanos S et al. 2011), to deliver thousands of short VNS trains in 2 freely behaving monkeys, over 11-16 hours, while recording intracortical LFPs across sites in prefrontal, sensorimotor and parietal areas. We identified epochs in which LFP activity in wide-spread cortical areas was dominated by high frequency oscillations (8-55 Hz), with and without overt movement, indicative of wakefulness, by theta oscillations (4-8 Hz), indicative of rapid-eye-movement (REM) sleep and by delta oscillations (1-4 Hz), indicative of late stages of non-REM (NREM) sleep. We compiled VEPs in each cortical site, separately for each of those states. We also documented the effects of pulsing frequency (5-300 Hz) on cortical responses. We found that VNS elicited VEPs in all sampled cortical areas. Most VEPs comprised 3 main components, with short (<70 ms), intermediate (70-250 ms) and long latencies (>250 ms). The magnitude of the early component was not affected by brain state, while that of intermediate and late components was significantly larger during sleep. At the same time, pulsing frequency of VNS significantly affected the amplitude of VEPs. These effects were sizable, as VEPs elicited from 300-Hz VNS during epochs of NREM sleep had intermediate and late components that were 300-500% larger than those during awake conditions.

## Methods

### Subjects

Experiments were performed with two male macaque (nemestrina) monkeys aged 5 and 6 years old and weighing 10.4 and 10.9 kg, respectively. The experiments were approved by the Institutional Animal Care and Use Committee (IACUC) at the University of Washington and all procedures conformed to the National Institutes of Health Guide for the Care and Use of Laboratory Animals.

### Cortical implant

During sterile surgery, each monkey was anesthetized with sevoflurane gas. A midline scalp incision was made and the scalp was resected. The intracortical electrodes were implanted through individual 0.5mm burr holes drilled with a stereotaxic guide. A total of 32 electrodes were placed in two hemispheres. On each hemisphere, the electrodes were located over the prefrontal, sensorimotor and parietal cortical areas. M1 received also two penetrating electrodes targeting the thalamus (one for each hemisphere).

The intracortical electrodes were made in house with 3mm cut length of platinum rod (AM Systems #711000, 0.254 mm diameter) insulated with heat-shrink Pebax (Small Parts #B0013HMWJQ). Pebax was cut so that the diameter of the exposed tip was ∼0.5 mm, corresponding to an exposed surface area of ∼0.06 mm^2^. Impedances ranged between 10 and 50 KOhms (at 1000 Hz). Skull screws used as ground and reference leads were placed on the occipital or the temporal bone, depending on skull exposure during surgery and the availability of space after the electrode implantation. The implant and the connectors were secured to the skull with acrylic cement and enclosed in titanium casing that was also attached to the skull with cement and skull screws.

### Vagus nerve implant

In a separate procedure the monkeys received the stimulating cuff on the left cervical VN. Under general anesthesia with sevoflurane gas each animal was positioned supine. A horizontal cervical incision was made above the left clavicle, medial to the sternocleidomastoid muscle. The skin was retracted to obtain a vertical exposure of the carotid sheath. The sheath was then opened, and the VN was exposed for a length of ∼3-4 cm between the jugular vein and the carotid artery. A rubber loop was passed behind the nerve to help with the manipulation and the VN electrode was placed around the VN trunk. We then secured the proximal end of the electrode leads with sutures at the subcutaneous tissue close to the implant to provide support for the placement of the nerve cuff on the nerve trunk. The distal end of the leads was routed subcutaneously all the way up to a skin opening at the back of the head, very close to the edge of the head chamber that has been installed previously during the cortical procedure (see skull surgery and cortical implant). The leads immediately entered the chamber and they were secured to the base using acrylic. A two-channel connector was used to electrically connect the cuff leads with a stimulator. The head and the neck incisions were then sutured. The monkey was treated with post-surgery analgesics and antibiotics for a 10-day recovery period.

A bipolar nerve cuff (Cyberonics, LivaNova Inc.) comprised two platinum-iridium filament bands each of them embodied in a flexible, silicone-base, helical loop. A third helical loop was included to stabilize the array around the nerve. The total length of the cuff (3 helical loops) measured 3cm with an internal diameter of 2mm.

### Overnight recording and stimulation

We used an updated version of Neurochip2 (Zanos S *et al*. 2011), the Neurochip3, to record cortical activity and motor movements while simultaneously stimulating the vagus nerve, for a total period of 10-16 hours. Signals from 16 cortical sites (maximum number of Neurochip3 recording channels) from the right hemisphere, contralateral to the implanted nerve cuff, were recorded single-ended, with a skull screw as tissue ground, at 16-bit resolution, sampled at 5 KHz, and a low-frequency cutoff of 0.01Hz. The choice of recording neural signals only from contralateral sites was dictated by our earlier observation that VEPs in the contralateral hemisphere were overall larger than those in the ipsilateral hemisphere (Zanos S, Moorjani S., Sabesan S., Fetz EE 2016). Gross motor movements (head and whole-body movements) were quantified by a 3-axis accelerometer powered by a 3 V lithium coin cell fixed in the titanium casing. The three analog outputs of the accelerometer were passed through a sum-of-absolute circuit and the magnitude of its signal output was sampled at 5 KHz.

Neurochip3 is also equipped with a stimulator specifically designed to meet the need for minimum power consumption, low component volume, high voltage capability (compliance to at least ±50V), wide current output range (accurate from 10μA to more than 5mA), true biphasic capability, and high output impedance. Thanks to these features we reliably delivered trains of stimulus pulses in current mode through a bipolar stimulation channel connected to the nerve cuff leads (see next paragraph for details). The impedance value between the two cuff contacts was in the range of 5 KOhm after implantation. Considering a current intensity between 1.0 mA and 1.5 mA, the returned voltage output was between 5V and 7.5 V, safely below the voltage limit of the stimulator. Stimulus timestamps were stored in the same time-base as the neural recordings.

Each recording began with the animal seated in a primate chair in the lab. Neurochip3 was then programmed by entering the desired settings into a Matlab GUI and uploading them via IR connection. The animal was then returned to its cage where it moved freely until the following day. Recorded data were stored on a removable flash memory card with 32-GB capacity and later imported to Matlab. For each session we took notes of the time of day when the Neurochip started to record.

### Vagus nerve stimulation

The Neurochip was programmed to deliver trains of 200 μs biphasic, symmetric, current pulses at an intensity of 1250 µA (in a separate control experiment in M1 we also used 1000 µA and 1500 µA). Each train consisted of 5 pulses with 10-sec interval between consecutive trains. Neurochip3 cycled through 4 different pulsing frequencies during each session. The tested frequencies were arranged in two sets. The first set was defined as [5Hz, 30Hz, 100Hz, 300Hz], while the second set contained [50Hz, 80Hz, 150Hz, 200Hz] as frequency values. The two frequency sets together resulted in a total of 8 stimulation protocols, tested in multiple sessions of 4 stimulation protocols per each session. In each session the same stimulation frequency was delivered for 20 or 10 minutes, in M1 and M2, respectively, before switching to the next stimulation frequency. The 4 stimulation frequencies kept cycling without breaks until the batteries completely discharged (on average 12 hours). Given these settings, the subject received 60 trains of stimuli (300 pulses total) at a given stimulation frequency every 10 minutes of recording. Thus, in a representative recording of 12 hours long the subject received around 4,320 trains of 5 VNS pulses without breaks. We completed 4 sessions with M2, with the 8 stimulation protocols tested twice, and 2 sessions with M1 resulting in a total number of 7 stimulation protocols tested once (the 30Hz stimulation was not delivered, data without stimulation was retained instead).

### Brain state classification

Data analysis was performed offline in Matlab (MathWorks, Inc.) through customized scrips. The identification of brain states was performed by evaluating the contribution of different frequency bands on the power spectrum of the LFP signal for each cortical site and by estimating the amount of movement assessed through the acceleration signal. Brain state classification was performed on 6-sec epochs preceding the delivery of electrical stimuli (Figure S1A, https://doi.org/10.6084/m9.figshare.12724739.v6), and four different states were identified (Figure 1). AW state was attributed to epochs with acceleration above zero for a sustained period of time (>60% of the duration of the epoch). RW epochs were characterized by neural signal with a predominant contribution of alpha (8 – 14 Hz), beta (14 – 35 Hz), and gamma (35 – 55 Hz) activity. Epochs showing high delta activity (1 – 4 Hz) in the EEG were scored as NREM, while the REM state was associated to a predominant theta activity (4 – 8 Hz). All these three latter states were accompanied with acceleration kept below the threshold used to assess the AW state. Our classification criteria also took into account the time of day, by assigning the two sleep stages (NREM and REM) only to epochs between 6 pm and 7 am, times when the room light went off and on respectively. This last criterion was included only to refine the classification algorithm and did not compromise the final outcome. This is demonstrated by the comparison based on the number of classified epochs for each recording site and each brain state between the classifier which included the time of day as a classification criterion and the classifier which did not include it as a criterion. Both classifiers returned a similar number of epochs (Figure S2, https://doi.org/10.6084/m9.figshare.12724739.v6).

**Figure 1:**
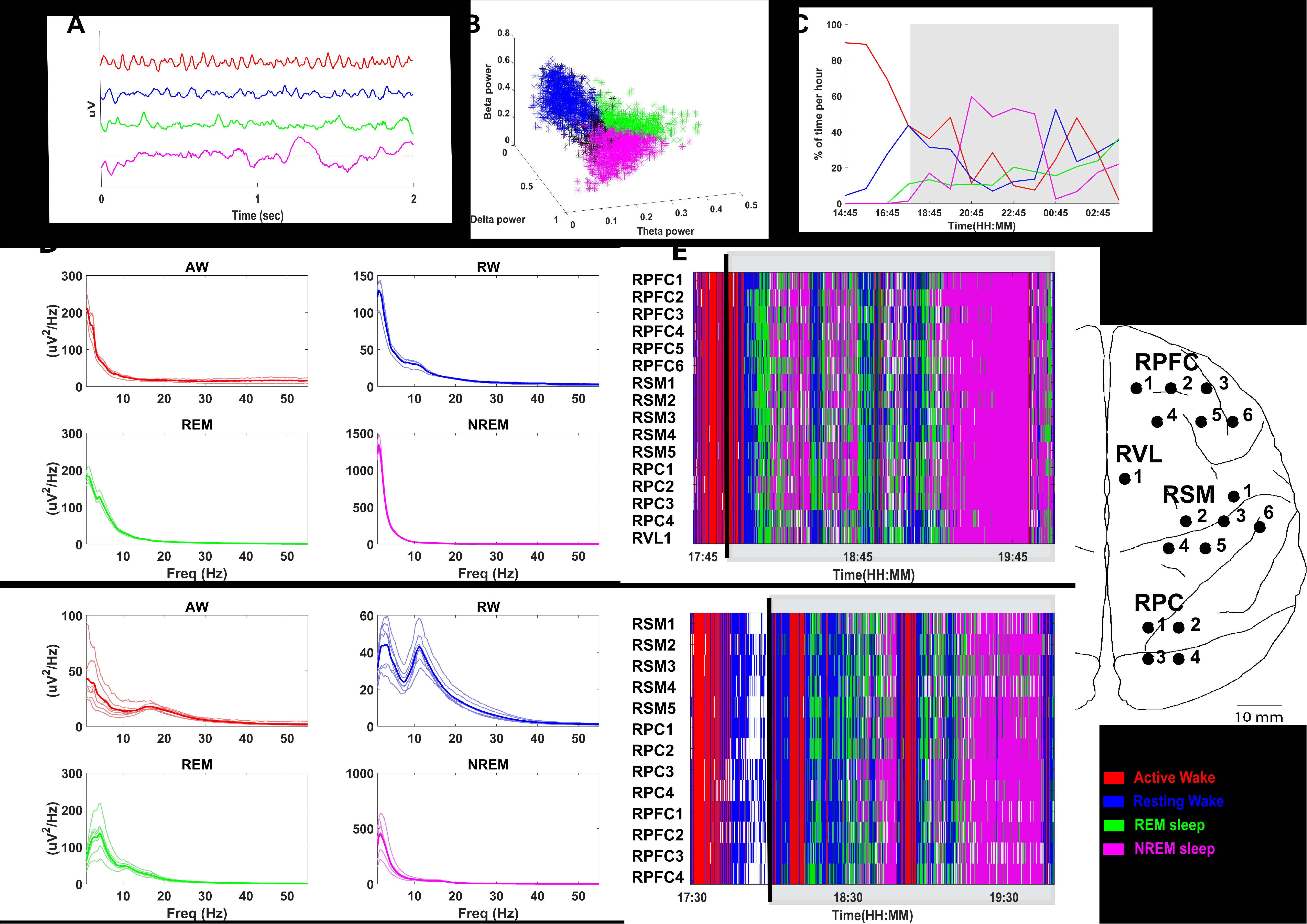
Different brain states using classification strategy based on the power in different frequency bands were successfully discriminated and the classified brain states showed a characteristic power spectrum profile with a global trend across recording sites for both animals. A) Two seconds of raw signal for each classified state of one representative cortical site from M2. B) Relative power in beta, delta and theta bands of classified epochs throughout the recording of the same cortical site shown in A (in black are shown the unclassified epochs). C) Percentage of time per hour spent in each brain state as a function of time of day. The gray-shaded area indicates the time during which lights were off. D) Power spectrum profile of classified epochs over all cortical channels for each recording (thin traces). Thick traces show the average across the recordings (M1, top rows; M2, bottom rows). E) Classified epochs for each cortical site over two hours of one representative recording. The gray-shaded area indicates the time during which lights were off (M1, top rows; M2, bottom rows). The location of the recording sites is represented on the right. RPFC: right prefrontal cortical area; RSM: right sensorimotor cortical area; RPC: right parietal cortical area; RVL: right ventral lateral caudal nucleus (thalamus). Red: active-wake; blue: resting-wake; green: REM, pink: NREM. Unclassified epochs are represented in white.

### Power analysis

Neural recordings from each cortical site were segmented into 6 second-long epochs, taken before the onset of each VNS train (Figure S1A, https://doi.org/10.6084/m9.figshare.12724739.v6) delivered every 10 seconds throughout the experiment duration. For each epoch we calculated the LFP power spectrum using the multi-taper method (Babadi B and EN Brown 2014) and we estimated the absolute power in several frequency bands: delta 1 – 4 Hz, theta 4 – 8 Hz, alpha 8 – 14 Hz, beta 14 – 35 Hz, gamma 35 – 55 Hz, as the integral of the absolute power density (uV^2^/Hz) within each frequency range (Figure S1A, S1B, https://doi.org/10.6084/m9.figshare.12724739.v6). We then derived the relative power in each band by dividing the absolute power by the total absolute power summed over all 5 frequency bands (Figure S1B, https://doi.org/10.6084/m9.figshare.12724739.v6). For each channel, we obtained 5 distributions of relative power values, with the same number of samples (i.e. the total number of epochs during the duration of each free behavior experiment). The distributions of relative power values were then log-transformed (log(x/1-x)), to convert them to Gaussian (Gasser T et al. 1982) (Figure S1D, https://doi.org/10.6084/m9.figshare.12724739.v6).

We then estimated the variable z by normalizing the Gaussian relative power distribution using z-score normalization:

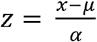, where µ is the recording mean of the transformed variable x and α is its standard deviation (Figure S1D, https://doi.org/10.6084/m9.figshare.12724739.v6).

### Detection of movement

The acceleration signal was smoothed with a moving average filter of 10 msec duration. A positive threshold was applied to the resulting signal in order to assess when the monkey was moving. The filtered signal had to remain continuously above threshold for at least 300 ms to be considered as “movement”; if the time lag between two consecutive movements was less than 3 sec they were merged together and treated as a single movement. The outcome of this process was a binary state vector of the same size as the original acceleration signal with 1 denoting “animal moving” and 0 denoting “animal resting”. For each 6-sec epoch prior to stimulation onset, we assessed the amount of movement by calculating the percent of time the animal was moving during each epoch. Consequently, we assigned a number ranging from 0 to 100 which expressed the amount of movement for each epoch (e.g. 60 denotes that the animal moved for 60 percent of the total epoch duration).

### Classification of brain states

Throughout the duration of the experiment, for each 6-sec epoch, we obtained a set of 7 features:

1. Amount of movement (assessed by the acceleration signal)
2. Room light ON and OFF (assessed by the time of day of that epoch, i.e. light OFF between 6pm and 7am)
3. Z-scored distribution of relative power in delta band (delta-power)
4. Z-scored distribution of relative power in theta band (theta-power)
5. Z-scored distribution of relative power in alpha band (alpha-power)
6. Z-scored distribution of relative power in beta band (beta-power)
7. Z-scored distribution of relative power in gamma band (gamma-power)

The algorithm for classification of brain states relied on the presence of movement and the contribution of different power bands to the cortical signal relative to a threshold value, which defined what was considered “high” or “low” power in all bands. Each epoch was then assigned to one of four brain states using a set of criteria:

- Active-wake (AW): Presence of movement for more than 60% of the duration of an epoch
- Resting-wake (RW): Movement for less than 60% of the duration of an epoch, high alpha-power (i.e. greater than threshold value), high beta-power, high gamma-power, low relative theta power (i.e. smaller than threshold value), low delta-power
- REM sleep: Light off, movement for less than 60% of epoch duration, high theta-power, low delta-power
- NREM sleep: Light off, movement for less than 60% of epoch duration, high delta-power

Because of the Z-score transformation, the relative power in each frequency band was scaled such that all bands contributed with the same “weight”. This way we were able to define a unique threshold power value for all frequency bands. This method ensured that multiple states were not assigned to the same epoch; at the same time, if none of the criteria was met, that epoch was considered “unclassified”. To find the optimal threshold value that discriminated the four brain states, we calculated the number of unclassified epochs for each recording site as a function of different Z-score threshold values, ranging from −3 to 3 in steps of 0.1 (Figure S1C, https://doi.org/10.6084/m9.figshare.12724739.v6). We then used the threshold value which minimized the number of unclassified epochs (Figure S1C, S1D, https://doi.org/10.6084/m9.figshare.12724739.v6).

The choice of 60% (of the duration of the epoch) as the threshold for considering an epoch as a “movement” epoch was dictated by several considerations. To assess when the animal was moving we used the acceleration signal. We merged two consecutive movements when the time interval between the two was less than 3 s. The movement threshold of 60% was chosen because it corresponded to a reasonable time period (3.6 s) consistent with the criteria used to analyze the acceleration signal. In animal M2 a movement threshold of 60% returned a balanced number of AW epochs versus RW epochs (Supplementary Figure S3 (https://doi.org/10.6084/m9.figshare.12724739.v6). In animal M1 the threshold which returned the same number of AW and RW epochs was 85%. The morphology of the classified VEPs did not show significant differences when the movement threshold was set at 85% vs. https://doi.org/10.6084/m9.figshare.12724739.v6).

### Compiling of vagal-evoked potentials

A stimulus-triggered average of LFP activity in each recording site was compiled to produce the vagal-evoked potential (VEP) at that site. The stimulation artifact was suppressed by linear interpolation between single voltage samples 0.2 ms before and 2 ms after the onset of each pulse in a VNS train. To assess how brain state modulated VEPs, each LFP trace from 100 ms before to 900 ms after the first stimulus in a train was assigned to the brain state classified from the 6-sec epoch that preceded the onset of the corresponding stimulus train. VEPs associated with a given state were then compiled by averaging all LFP traces assigned to that state. Since there were multiple sleep cycles during a given 10-16 hour-long experiment, single traces assigned to a given state were not necessarily recorded during the same cycle.

VEPs in each recording site were characterized by the amplitude of the biggest deflection (either positive or negative) within each of 3 VEP components. Each VEP component was defined by a latency window, measured from the first pulse in the stimulation train: an early component between 5ms and 70ms, an intermediate component between 70ms and 250ms, and a late component between 250ms and 600ms. The 3 latency windows were defined empirically and with practical considerations in mind, given the variability of VEP shapes across animals, brain states and cortical sites. With the 5-70ms latency window, we attempted to capture the early VEP component that remained relatively invariant with respect to brain state. Typically, this was a single, monophasic or biphasic waveform. VEP deflections in the 70-250 ms latency window were quite variable between brain states and animals. Subdividing this window into more than one windows did not offer any more insights into the modulation of VEPs by brain state and quantifying the amplitude of individual deflections in this range provided several VEP measures that had little consistency between animals and brain states. The neurophysiological mechanisms underlying each of those deflections was also unclear. For those reasons, we decided to characterize VEP deflections within that latency window using a single component. The downside of this approach is that the amplitude measurement in this window is sensitive to relative changes in the amplitude of individual waves, which may explain some of the polarity changes we observed (Figure 5). Finally, the portion of the VEP with longer latencies, typically 250-600 ms, almost always comprised a single, slow negative deflection, large in amplitude, modulated strongly by brain state.

In addition to amplitude of the largest deflection, we considered other measurements for each of the 3 VEP components, including peak-to-trough amplitude and root-mean-square (RMS). The RMS is an approximation of the area under the curve and is affected by the amplitude (maximum deflection) but also by the duration of the non-zero components of the VEP. These measurements returned comparable results with respect to brain state modulation.

In order to quantify the variability of the individual evoked responses for a given brain state, we estimated the single-trial amplitude by taking the inner product of a single trial trace with the normalized VEP associated with the same brain state as that trial. More specifically, the VEP associated with a brain state *S* was defined as:

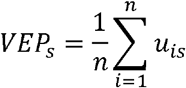

where u_is_ (i=1,2,..,n) were all the single trial traces classified into brain state *S.* By normalizing the VEP_s_ by the square-root of the average power we obtained the “template waveform” for brain state *S*:

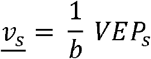

Where:

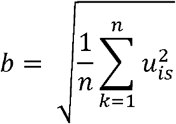

This in turn was used to calculate the single-trial amplitude *A_is_*, taken as the inner product between a single trial trace classified into brain state *S* and the template waveform for that brain state:

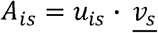

This approach takes into account the shape and the size of the entire waveform and the single trial amplitude is determined not only the larger deflection, but also from the degree of similarity to the average waveform. This measurement provided a direct comparison of the single trial variability among brain states.

### Coherence and phase estimation

To estimate the effects of VNS on cortical synchrony we calculated the pairwise coherence spectrum for each classified epoch for all electrode pairs using LFP signals recorded during two time windows with respect to the onset of the stimulus train: −3 to −1 s (“pre-VNS”) and 2 to 4 s (“post-VNS”). The choice of these time windows allowed us to assess VNS-elicited changes in coherence between cortical sites, from pre-stimulation baseline, without the confounding effect of stimulus-evoked activity appearing on many cortical sites simultaneously and therefore “artificially” increasing coherence. After down sampling the signals to 500Hz, coherence was compiled in the time-frequency domain as magnitude-squared wavelet coherence (Romcy-Pereira RN et al. 2008; Qassim YT et al. 2013) for both time windows. The change in coherence, between post-VNS and pre-VNS, was then calculated as the difference of mean coherence within each frequency band. This calculation was performed separately for each epoch; because of the large number of epochs, we used t-test to assess the significance of these changes in coherence.

In order to assess whether the phase of delta oscillatory activity affects the modulation of VEP amplitude by state, we extracted the analytic phase in the delta band from epochs classified in the NREM state (the brain state with high delta power), using Hilbert transform. We then compiled VEPs by separately averaging those traces in which stimulus onset occurred during a hyperpolarizing phase of delta (phase angle [0-π] rad), and those traces in which stimulus onset occurred during the depolarizing phase of delta (phase angle [π-2 π] rad). To compare the 2 groups of VEPs, we calculated the root-mean-squared (rms) for the whole response window of 5-600 msec from stimulation onset for each recording site.

### Statistical analysis

To assess whether VEP modulation by brain state modulation was affected by VNS parameters we compiled separately VEPs for each set of parameters (“VNS protocols”). Three-way analysis of variance was used in order to test for statistically significant differences in the VEP amplitude between brain states, VNS protocols and cortical areas. We also performed a post-hoc, pairwise comparison test of means, with Bonferroni correction, between brain states.

### Median nerve stimulation

To test whether the effect of brain state is specific to VNS and VEPs, we performed control experiments in which the median nerve was stimulated through an electrode pair placed on the skin of the anterior forearm, to evoke somatosentory evoked potentials (SSEPs). To protect the skin electrodes from the monkey’s reach, we performed these experiments in the booth, with the animal in the primate chair and its arms restrained. To facilitate the animal falling asleep, these experiments were performed in the evening, in quiet conditions, with the booth lights turned off. Out of three tested monkeys, only one (M3) was able to relax and eventually fell asleep, as confirmed from an infrared camera. At times, the animal woke up, opened its eyes, and moved its body and legs. It is therefore unlikely that the animal was able to cycle through different sleep stages, as was the case in unrestrained conditions. With that in mind, we modified the classification of brain states. We scored the 6 s epochs based on the prevalence of delta oscillations: from epochs showing a minimum of delta power contribution (D1) to epochs mostly characterized by power in delta range (D4) (Figure S7 A, B, C, https://doi.org/10.6084/m9.figshare.12724739.v6). With this modified brain state classification (Figure S7C, https://doi.org/10.6084/m9.figshare.12724739.v6), states D1 and D2 are mostly characterized by increased power in alpha, beta and gamma ranges, indicative of wakefulness, whereas states D3 and D4 are mostly characterized by low frequency components.

## Results

### Classification of brain states

Using the power spectrum of the cortical signal, the accelerometer signal and the time of day, four different brain states were discriminated: active wake (AW), resting wake (RW), REM sleep and NREM sleep (Figure 1). The AW state, defined as acceleration value above zero for most of the duration of the 6 seconds epoch, occurred predominant during daytime (before 6pm, at which time housing room lights were turned off) with some brief periods during nighttime (Figure 1C); the associated cortical signal was characterized by more power in beta and gamma frequencies compared to the other three states (Figure 1A, 1D). RW, REM and NREM were defined by different relative contributions of five frequency bands to the total cortical power in the frequencies between [1-55] Hz. RW showed higher contribution of alpha frequency band which is typically associated to a resting state (Pfurtscheller G et al. 1996) (Figure 1A, 1D). REM sleep was characterized by more power in theta band than in any other frequency band, while NREM sleep was characterized by high amplitude oscillations in delta band (Figure 1A, 1D). The percent of time spent in NREM sleep decreased as the night progressed for both animals (Figure 1C, Figure S4, https://doi.org/10.6084/m9.figshare.12724739.v6), which is typical for non-human primates (Daley JT et al. 2006; Rachalski A et al. 2014). Although we assigned brain states at a much finer timescale than what has been previously used in sleep staging studies in monkeys (Hsieh KC et al. 2008) (6 s-long epochs vs. 10-30 sec-long epochs), we observed similar overall patterns: REM episodes clustered in time and were generally brief (3-10 min in duration), whereas NREM episodes were longer (20-40 min long) (Figure 1E). Even though our brain state classification was performed for each cortical site independently, most cortical sites “reported” the same brain state in a given epoch (Figure 1E). Requiring that all cortical sites report the same brain state in order for that epoch to be assigned to that brain state, had little effect on the sequence of epochs assigned to awake states, moderate effect on that of NREM sleep states and significant effect on that of REM sleep states (Figure S4B, https://doi.org/10.6084/m9.figshare.12724739.v6). Delivery of VNS did not affect the structure of brain states as the global number of transitions between states was not altered by the presence of stimulation itself nor by any specific VNS protocol (Figure S5, https://doi.org/10.6084/m9.figshare.12724739.v6). Because the brain state classification was based on power contributions of different frequency bands, this indicated that VNS delivered in trains did not overall have short-term effects on the power of specific frequency bands.

### Effect of brain state on VEP responses

Cortical VEPs elicited by delivering trains of electrical stimuli to the vagus nerve were characterized by three main components, early, intermediate and late, whose polarity and magnitude were dependent on cortical area and brain state; there were also differences between the two subjects (Figure 2). The 2 later VEP components were strongly modulated by brain state in both animals, especially at higher VNS pulsing frequencies (p<0.001 for both components and both animals in 3-way ANOVA for maximum deflection magnitude and RMS) (Figure 3, 4, 5). The early component did not show significant dependence on brain state, taken either as maximum deflection amplitude (p=0.47 for M1, p=0.58 for M2 in 3-way ANOVA) or RMS value (p=0.2 for M1, p=0.17 for M2). In contrast, intermediate and late components of VEPs had larger magnitudes during sleep than during wakefulness. Both components were larger during REM sleep compared to either of the two awake states. In M1, that difference was significant for both components (p<0.001 pairwise comparisons Bonferroni corrected, for both max deflection magnitude and RMS). In M2, there was a significant difference in both magnitude and RMS during REM sleep compared to both awake states for only the late component (p<0.001, Bonferroni corrected), while the intermediate component was significantly different between the two awake states (p<0.05 for max deflection magnitude, p<0.001 for RMS), but not between resting state and REM sleep (p=0.6 for max deflection magnitude, p=1 for RMS, pairwise comparison Bonferroni corrected). The difference in the late component between the two awake states was not significant for either animal (p>0.05 for both max deflection amplitude and RMS). The 2 later components had the largest magnitude during NREM sleep (Figure 2, 3, 4): at 300Hz pulsing frequency, the intermediate component was 452% and 189% larger and the late component was 566% and 409% larger during NREM than awake in M1 and M2, respectively. This brain state modulation was independent of VNS current intensity, at least in animal M1 in which three current intensities were tested (1000, 1250 and 1500 µA), with the same pattern of brain state modulation in all VEP components (Figure S6, https://doi.org/10.6084/m9.figshare.12724739.v6).

**Figure 2:**
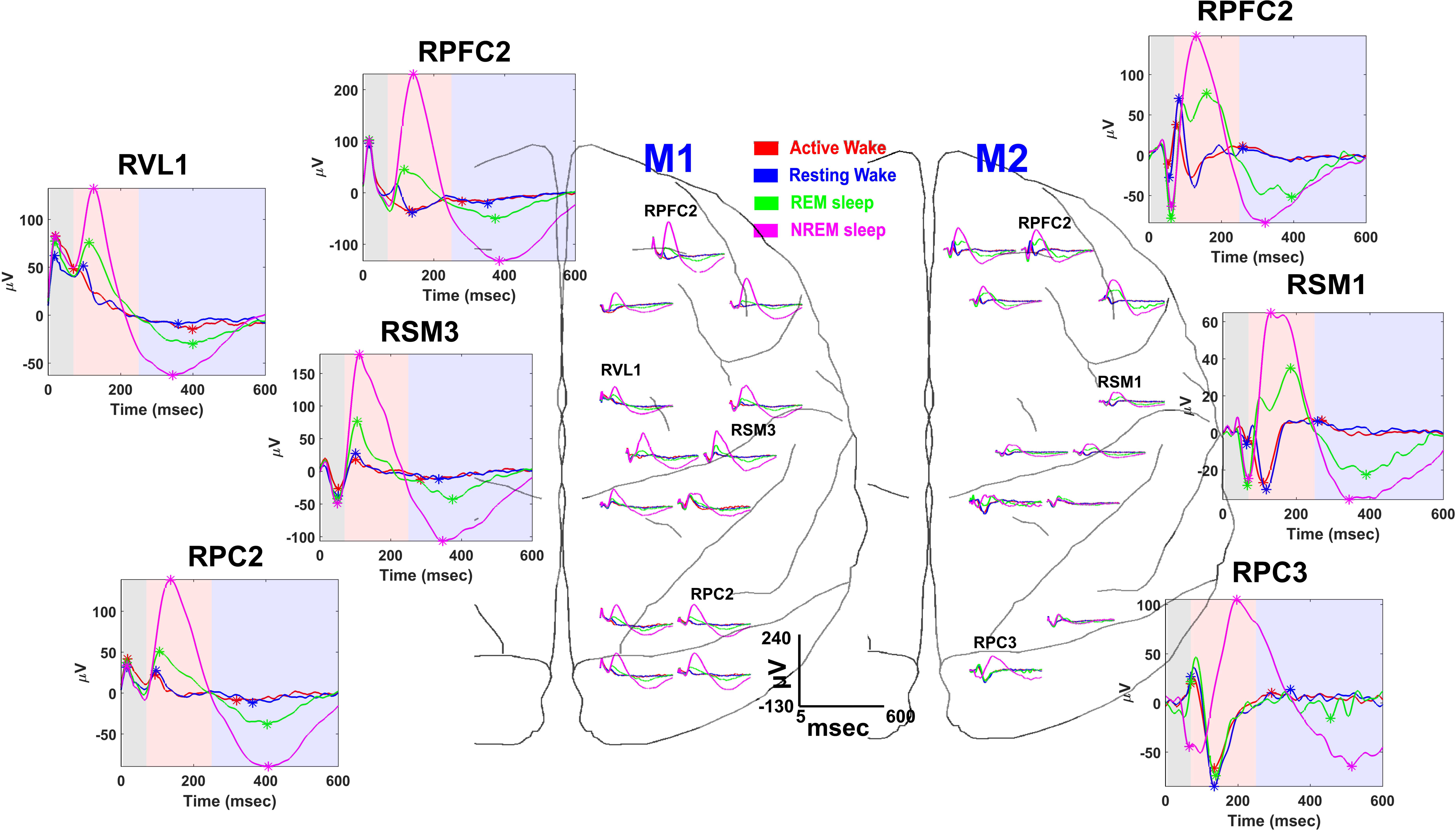
Vagal evoked potentials (VEPs) elicited during different states for both animals. VEPs were evoked by trains of 5 pulses at 300 Hz. Some representative recording sites for each animal are shown in the larger plots on the sides. The colored shadow areas highlight the time range for each VEP component: early, 5-70ms (gray); intermediate, 70-250ms (orange); late, 250-600ms (light blue). The detected components for each brain state are indicated by a colored *. The X axis represents the time after the first pulse in the train. The colored traces represent different brain states: red, active-wake; blue, resting-wake; green, REM; pink, NREM.

**Figure 3:**
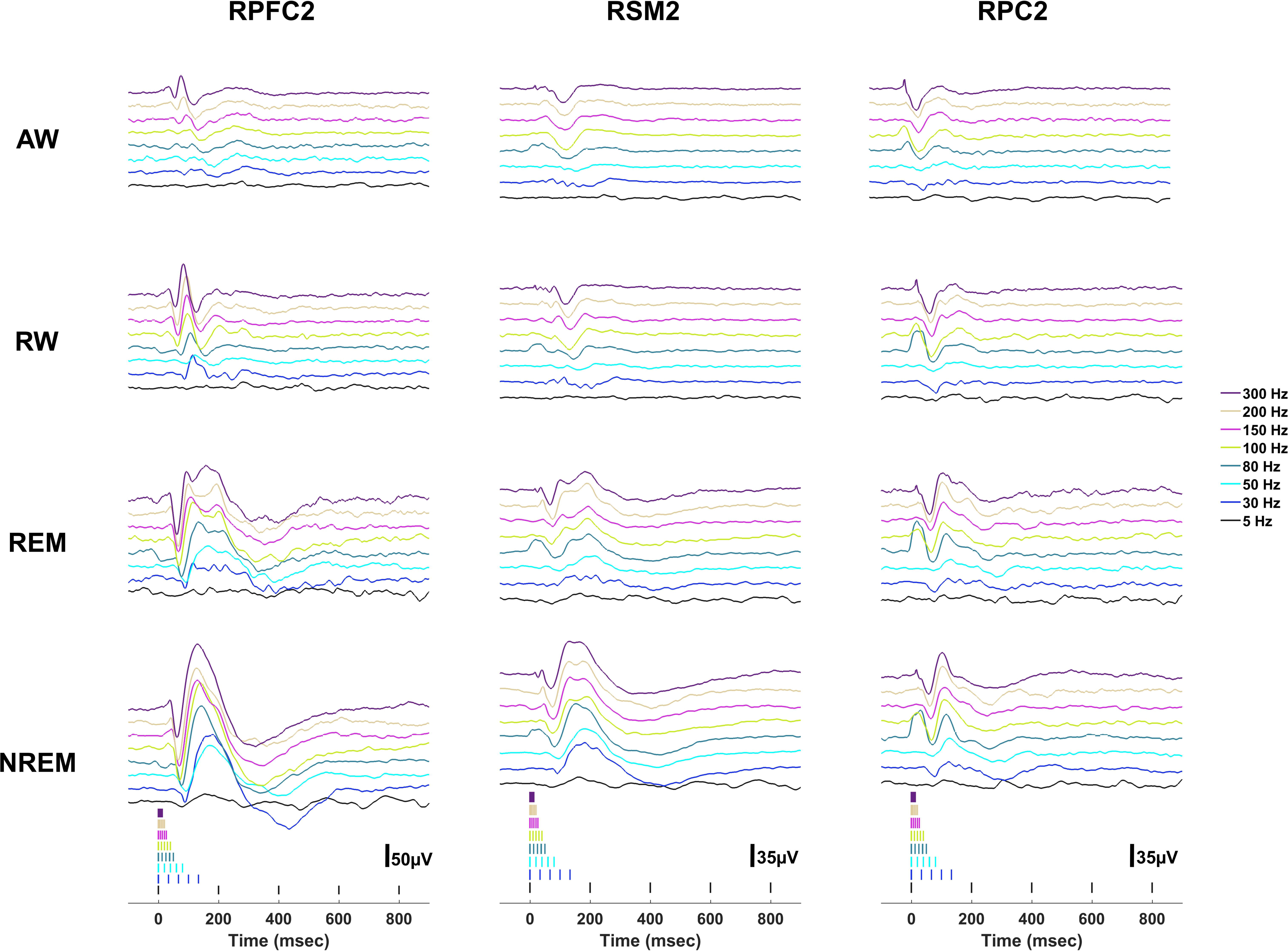
Vagal evoked potentials (VEPs) elicited during different states recorded from three representative sites of three different cortical areas (from left to right: prefrontal, sensorimotor, parietal) of animal M2. Colored lines show VEPs evoked by trains of 5 pulses at different pulse frequencies. VNS pulse times are shown below. AW: active-wake; RW: resting-wake; REM: rapid eye movement sleep; NREM: non-rapid eye movement sleep.

**Figure 4:**
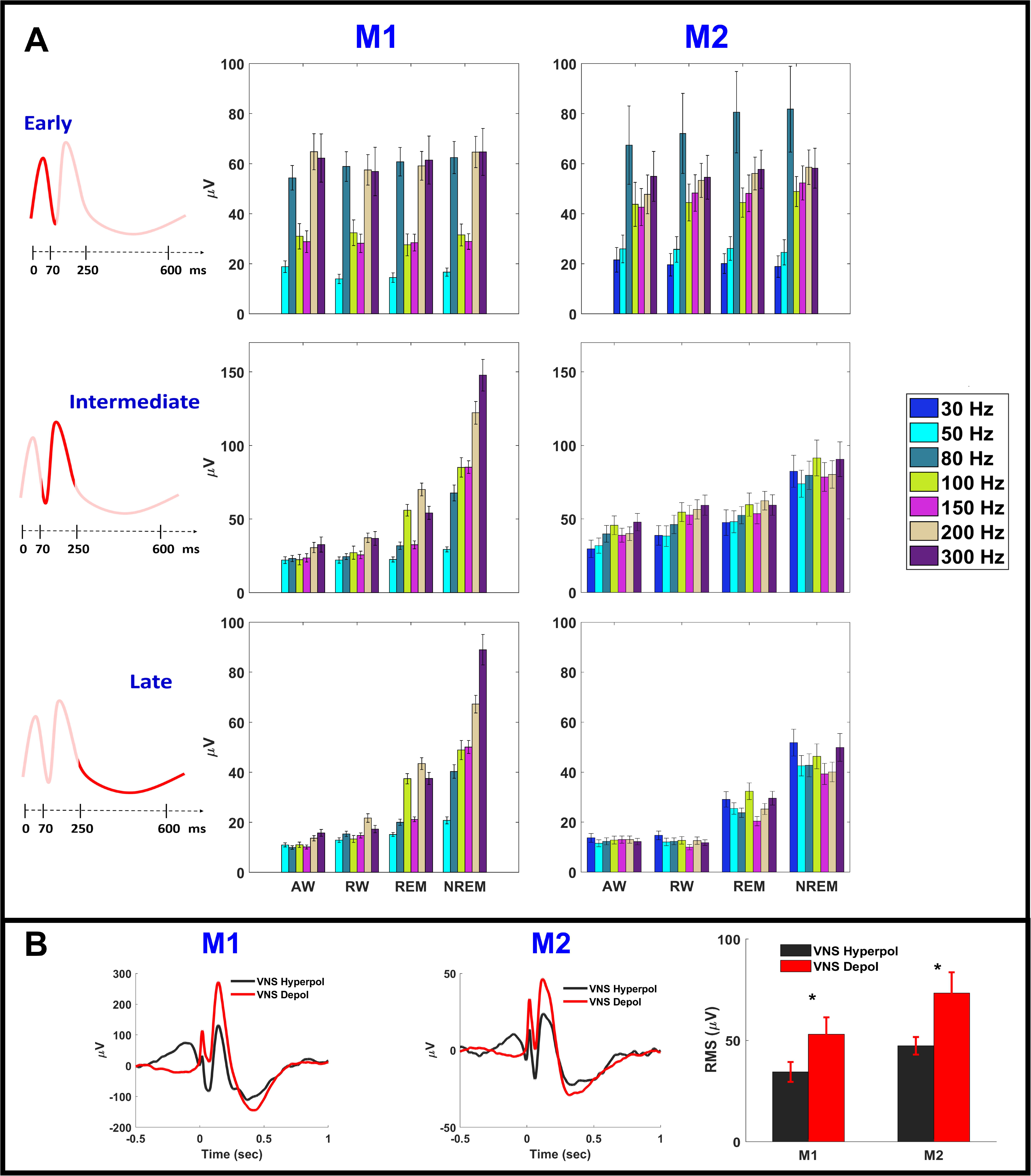
Modulation of VEP components by brain state, stimulation frequency and phase of oscillatory activity in delta range for both animals (M1, M2). A) Each group of bars represents the averaged absolute value of the maximum deflection of the VEP component over all channels (mean ± SE) evoked by trains of 5 pulses delivered at different stimulation frequencies (color coded) as function of brain state (X axis). Each VEP component (from top to bottom: early, intermediate, and late) was defined in a specific time window from the first pulse of the stimulation train (5-70ms, 70-250ms and 250-600ms, respectively). Both animals returned significant differences in VEP peak amplitudes between brain states for both intermediate and late component (n-way ANOVA: Intermediate, p<0.001; Late, p<0.001). The early component showed no significant difference (n-way ANOVA: Early M1, p = 0.766; Early M2, p = 0.848). All three components showed significant differences across stimulation frequencies (n-way ANOVA, p<0.001). B) Delta phase modulation of VEPs for both animals (M1, M2) during NREM. VEPs from one representative electrode for each animal evoked by 300Hz VNS during NREM. The traces are categorized based on the phase of delta oscillation at the stimulation onset: VEPs elicited by VNS delivered at hyperpolarized phases (0-π) are shown in black, whereas in red are shown the VEPs elicited by VNS delivered at depolarized phases (π-2π). The plot on the right shows the averaged root-mean-square (RMS) values of VEPs across all channels (mean ± std) calculated over a response window of [5-600] msec from stimulation onset. In average the VEPs elicited by VNS delivered at the depolarized phases of delta oscillations are larger than the ones elicited by VNS delivered at the hyperpolarized phases for both animals (ttest: p<0.05).

The same dependency of VEP magnitude on brain state was found when we considered the peak-to-trough measure for each of the three VEP components. This was expected, given the strong linear correlation between the amplitude of maximum deflection and the peak-to-trough magnitude (Pearson correlation coefficient R=0.94, p<0.001 for M1 and R=0.89, p<0.001 for M2) (Figure S8, https://doi.org/10.6084/m9.figshare.12724739.v6).

Importantly, the phase of ongoing delta oscillations affected the modulation of VEP magnitude by brain state (Figure 4B). During NREM sleep, VEPs elicited by VNS delivered at hyperpolarizing phases of delta cycles (0-π rad) were significantly smaller than VEPs elicited by VNS delivered at the depolarizing phases (π-2π rad), in both animals (Figure 4B).

After quantifying the variability in the shape of individual vagal-elicited responses across brain states, we found that during NREM sleep, the variability was minimal, in all cortical sites, and in both animals (Figure 5C). During REM sleep, only the prefrontal sites of animal M2, but not M1, had smaller VEP variability than awake states (Figure 5C).

**Figure 5:**
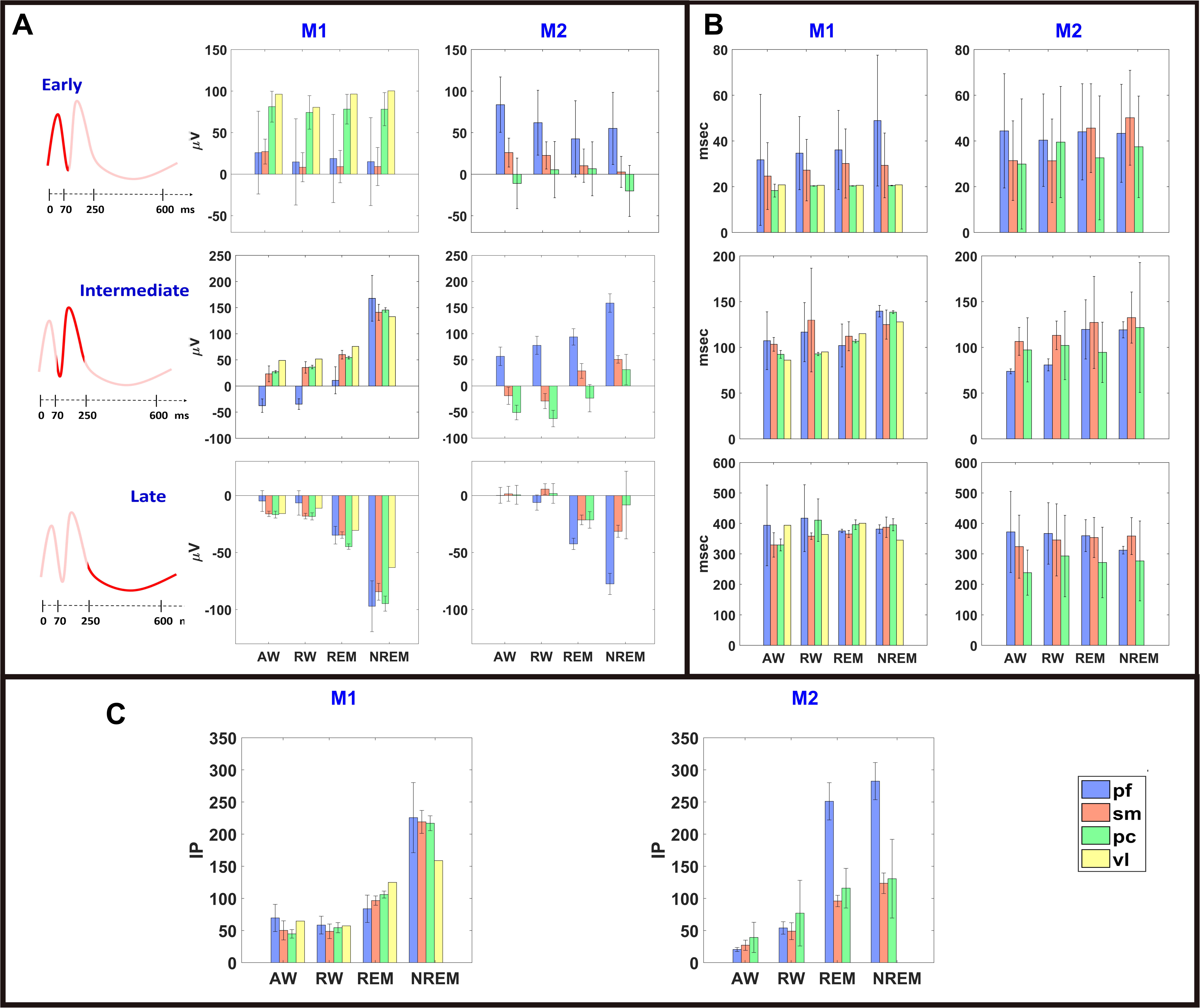
Brain state modulation of VEPs responses and VEPs variability for each cortical area for both animals (M1, M2). PF: prefrontal (blue); SM: sensorimotor (pink); PC: parietal cortical area (green); VL: ventral lateral caudal nucleus (yellow). VEPs in these plots were evoked by VNS trains at a pulse frequency of 300 Hz. A) Classified VEP magnitude (grand mean ± grand SE) for each component. B) Classified VEP latency (grand mean ± grand SE) for each component. C) Inner product (IP) between each classified single VEP sweep and the corresponding averaged VEP calculated between 5msec and 600msec from stimulation offset (grand mean ± grand SE over all channels for different cortical areas).

To assess whether lighting conditions affected the shape of VEPs independently of brain state, we compared VEPs classified to “awake” states during daytime (lights on) with VEPs classified to “awake” states during nighttime (lights off) (Figure S5B, https://doi.org/10.6084/m9.figshare.12724739.v6). The shapes of the VEPs were comparable between daytime and nighttime. This observation allowed to validate our choice to combine all the classified VEPs together, even if they were not necessarily recorded during the same sleep/awake cycle. Thus, we obtained a total number of epochs during awake states comparable with the total number of epochs scored as sleep (Figure S2A, https://doi.org/10.6084/m9.figshare.12724739.v6), even if most of the recordings happened during times of light off.

Finally, in a third animal (M3), wakefulness and sleep affected the shape of SSEPs elicited by median nerve stimulation, suggesting that modulation of evoked responses by brain states is not specific to vagus nerve stimulation (Figure S7, https://doi.org/10.6084/m9.figshare.12724739.v6).

### Effect of pulsing frequency on VEP responses

VEP responses were significantly modulated by pulsing frequency (p<0.001 for peak deflection and RMS for all three components in both monkeys, 3-way ANOVA). Higher stimulation frequencies elicited larger VEP responses in both monkeys, especially with regards to the intermediate and late components, whereas the effect on the early component was less consistent (Figure 4). Higher pulsing frequencies also evoked VEPs with a stronger modulation by brain state (Figure 4) and pulsing frequencies of at least 80Hz for M1 and 30Hz for M2 were necessary in order to evoke responses that varied with brain states (Figure 4).

We separately considered VEP responses elicited by VNS trains with a pulsing frequency of 5 Hz (Figures 3, 6). At this pulsing frequency, VEPs with 3 identifiable components were not always elicited (Figure 6). In some cases, each pulse in the train elicited a VEP with both early and intermediate components, and in other cases only the early component was evoked (Figure 6). Overall, VEPs looked different from the VEPs evoked by higher frequencies and only some of the cortical sites showed brain state modulation (Figure 6).

**Figure 6:**
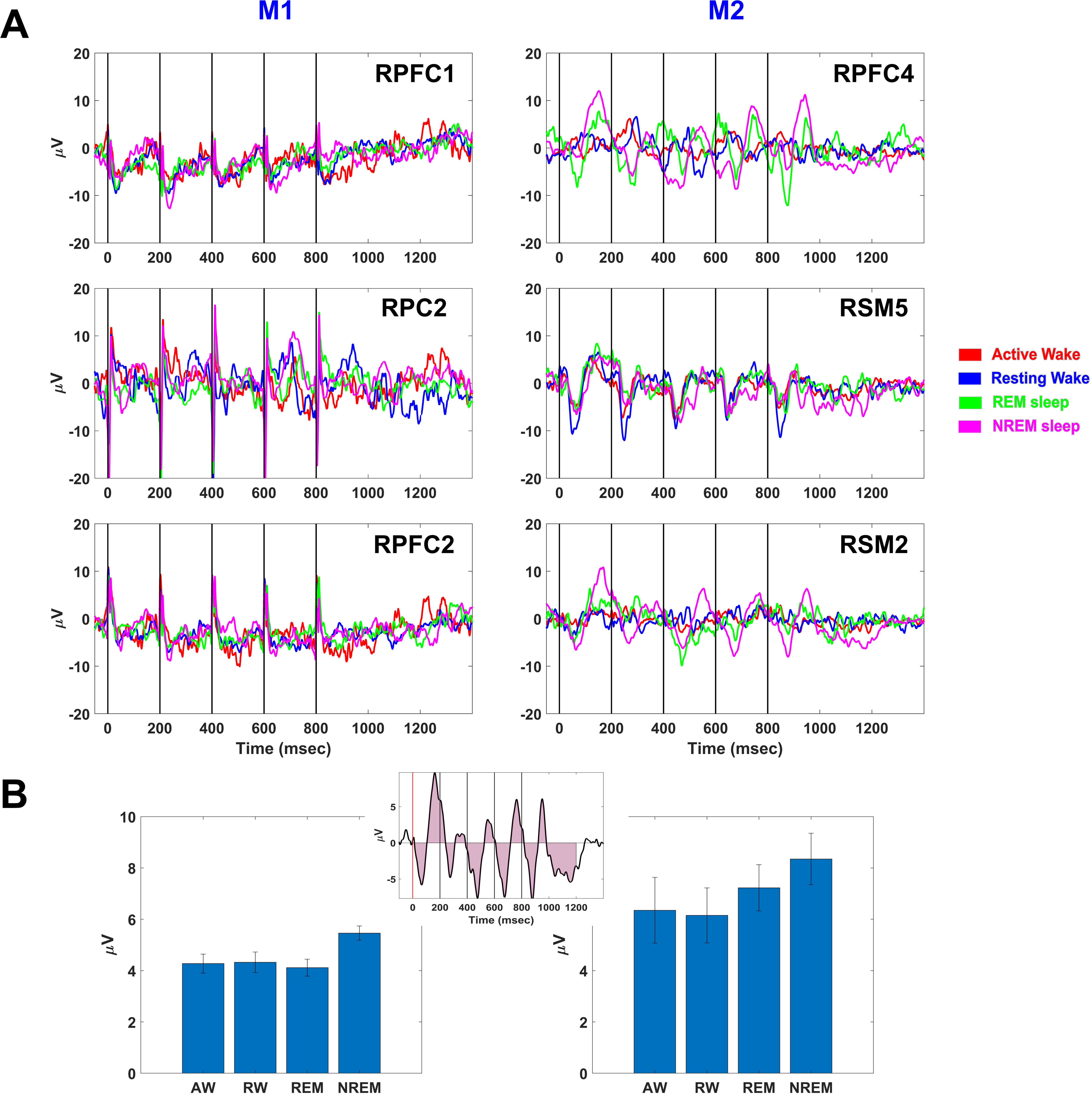
Classified VEPs evoked by 5 pulses VNS train at 5Hz in both animals (M1, M2). A) Example of classified VEPs for three different channels. The colored traces represent different brain states: red, active-wake; blue, resting-wake; green, REM; pink, NREM. B) Quantification of the classified VEPs as average over all channels (mean ± SE) of the responses’ root-mean square (rms) calculated between 10ms and 1200ms from stimulation onset. The inset on top shows a graphic representation of the rms measurement indicated by the pink area under the black curve.

Independently of their stimulation frequency, VNS trains did not show a significant (t-test, p<0.05) and consistent effect across the two animals on cortical coherence from pre-VNS levels, for any of the brain states (Figure 7, S10, https://doi.org/10.6084/m9.figshare.12724739.v6).

**Figure 7:**
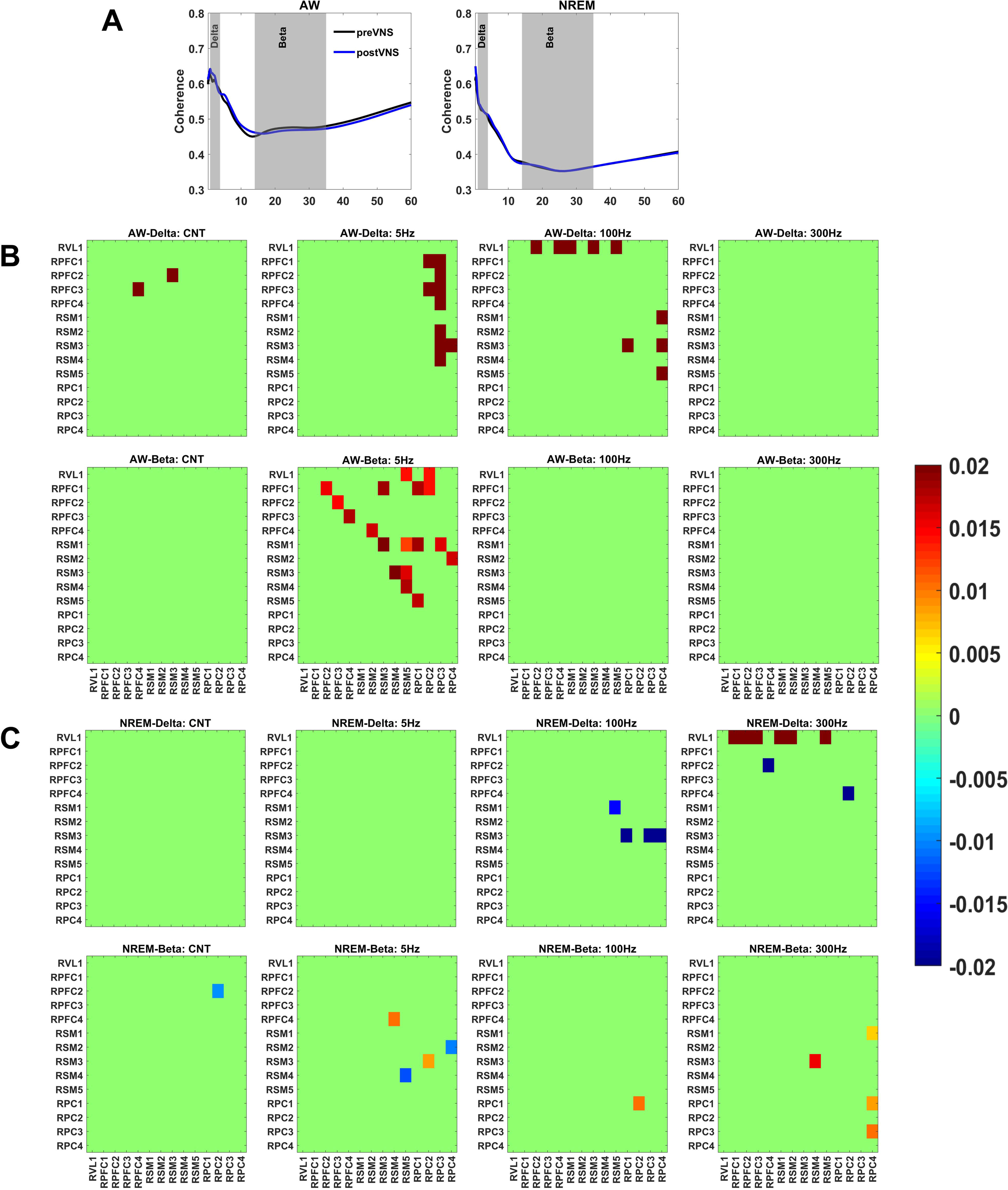
Effect of VNS on coherence for monkey M1. A) Coherence spectrum during awake (on the left) and during NREM sleep (on the right) calculated over two different time windows for one pair of channels: 2 seconds before stimulation onset (preVNS, black) and 2 seconds after VNS in a window between [2-4] sec from stimulation onset (postVNS, blue). The gray shadow areas highlight the two frequency bands considered to plot in the colormaps below the changes in coherence: beta band ([14-35]Hz) and delta band ([1-4]Hz). B) Color maps showing pairwise significant differences (t-test, p<0.05) in coherence in delta (top) and beta (bottom) range between all channels during awake for 4 different conditions: no stimulation delivered (CNT), 5Hz VNS, 100Hz VNS, 300Hz VNS. C) Color maps showing pairwise significant differences (t-test, p<0.05) in coherence in delta (top) and beta (bottom) range between all channels during NREM sleep for 4 different conditions: no stimulation delivered (CNT), 5Hz VNS, 100Hz VNS, 300Hz VNS. Refer to Figure 1 for the anatomical location of the electrodes.

### VEP responses in different cortical areas

To establish the dependency of VEP responses on cortical area, we compared the magnitude, polarity and latency of the main deflection of each VEP component elicited by 300Hz VNS across groups of cortical sites (prefrontal, sensorimotor, parietal and thalamic) for each brain state (Figure 5).

Early VEP component: In animal M1, the early component had small amplitude and variable shape at frontal and sensorimotor sites, while the responses at parietal and thalamic sites were positive and consistently larger (Figure 5A, 5B). In animal M2 the early component was positive at the prefrontal and sensorimotor sites and negative at the parietal sites, whereas prefrontal sites showed a consistently larger early component compared to other cortical areas (Figure 5A). Both animals had smaller early component amplitudes in sensorimotor sites compared to other cortical areas (Figure 5A). No obvious brain-state modulation of either magnitude or latency was observed for the early component.

Intermediate VEP component: In M1, the intermediate component was positive for all areas and brain states except at prefrontal sites, which showed a polarity switch from negative, during AW, RW and REM, to positive during NREM sleep (Figure 5A). The intermediate component for M2 was more variable across cortical areas. It was large and always positive in prefrontal sites, while in sensorimotor sites its magnitude was smaller and it switched polarity from negative during wake states to positive during sleep states (Figure 5A). The same component in parietal sites showed a polarity switch from negative during AW, RW and REM to positive during NREM sleep (Figure 5A). The peak latency did not show strong dependence on brain state in either animal (Figure 5B).

Late VEP component: In M1 the late component was negative with relatively consistent latencies across all cortical sites and brain states (Figure 5A, 5B). In M2 the amplitude of the late component was almost absent during awake states in all cortical areas, while during sleep states it was large, negative and with consistent latency (Figure 5A, 5B). Prefrontal sites in M2 showed larger amplitude compared to other cortical areas. In both animals the peak latency of this component was similar across cortical areas and was not modulated by brain state (Figure 5B).

## Discussion

### Brain-state modulation of VEP components

In this study we investigated the modulation by brain state of the cortical responses elicited by left cervical VNS in 2 freely behaving monkeys. To our knowledge, this is the first study to systematically document VEPs in nonhuman primates. The VEP responses were not identical between the 2 animals, even though similar cortical areas were sampled in both. That difference could be attributed to small differences in the way the cortical electrodes were fabricated and inserted into the brain, as well as in the exact location of the cortical electrodes, which was determined using non-subject-specific stereotaxic coordinates. It could also be related to the different implant age between the 2 subjects at the time of conducting these experiments (M1 had older implant than M2).

Four brain states were distinguished based on the power of different frequency bands in the local field potential (LFP) and the presence of movement, detected via a head chamber-mounted accelerometer: wakefulness in the presence of overt movement, wakefulness in the absence of movement, REM sleep and NREM sleep. In this study, LFP power was used as a marker to *infer* different brain states, usually defined with additional markers including electromyography and electro-oculography (EOG), which provides information on eye movements (Rechtschaffen, 1968). The EOG, for example, is critical to accurately defining the REM sleep stage that can be mistaken for the early stages of NREM sleep, as both are characterized by high power in theta range in the LFP (Armitage R 1995). In that sense, we cannot claim that the “brain states” used in the current study are necessarily the same as stages of sleep and wakefulness, only that ongoing cortical LFP activity during those times is indicative of those stages, as they share the same LFP criteria. Nonetheless, this approach is validated by several studies, in different species, which characterized the different awake and sleep states in term of power spectrum of cortical signal (Armitage R 1995; Rachalski A *et al*. 2014; Panagiotou M et al. 2017). Furthermore, other studies have shown that the most relevant feature for sleep/wake states classification is the set expressing the EEG relative power in different frequency bands, which alone is able to reach a classification accuracy above 70% (Zoubek L. CS, Lesecq S., Buguet A., Chapotot F. 2007; Charbonnier S et al. 2011; Krakovska A and K Mezeiova 2011). The accuracy increases even more when the EMG activity, here represented by the acceleration signal, is taken into account (Zoubek L. CS, Lesecq S., Buguet A., Chapotot F. 2007).

Moreover, in the originally defined criteria for sleep stage classification (Rechtschaffen, 1968) one electrode of the EEG montage, usually C3 or C4 in the international 10-20 system, is used. In our study, we treat cortical sites as independent from each other allowing the classification to be free from the assumption that all sites are in the same state at the same time. For example, the phenomenon of local sleep has been described in aquatic mammals, birds, rodents and humans (Vyazovskiy VV et al. 2011; Mascetti GG 2016). We found that the requirement that all cortical sites reported the same brain state before being assigned to that brain state, had minimal effect on the sequence of epochs with awake states, moderate effect on the sequence of epochs indicative of NREM sleep and significant effect on the sequence of epochs indicative of REM sleep (Figure S4B, https://doi.org/10.6084/m9.figshare.12724739.v6). Therefore, we find limited support for the assumption that when a “standard” EEG site (e.g. C3 or C4) reports a state, that state is representative of the states of the rest of cortical sites. This strategy allowed us to directly assess how LFP oscillatory dynamics at cortical sites, which correlates with brain states, modulated vagal evoked potentials at those sites.

We found that, independently of VNS intensity (Figure S6, https://doi.org/10.6084/m9.figshare.12724739.v6), brain state affected the magnitude of VEPs, especially of the 2 later components. Those 2 VEP components showed a progressive increase in magnitude from epochs indicative of wakefulness to those indicative of sleep, reaching maximum magnitude during delta dominant activity (NREM sleep) (Figure 4). It is unclear what cortical mechanisms are responsible for the generation of different components of VEPs, in sleep or awake states.

In Massimini et al. (Massimini M et al. 2007) cortical responses in the form of slow oscillations (SOs) were triggered in sleeping humans by transcranial magnetic stimulation (TMS) pulses. Slow oscillations (SOs) during NREM sleep represent the synchronous alternation between depolarized (“up-state”) and hyperpolarized (“down-state”) membrane potential of cortical neurons (Steriade M et al. 1993; McCormick DA and T Bal 1997; Destexhe A *et al*. 1999). In our study, the 2 later VEP components during sleep lasted approximately 500 ms, similar to the TMS-evoked responses in Massimini et al. (Massimini M *et al*. 2007) (Figure 2, 3). In both our and their (Massimini M *et al*. 2007) studies, evoked responses were state-, dose- and cortical site-dependent (Figure 4, 5). The prefrontal sites showed larger VEPs for M2 (Figure 5) and for M1 the magnitude of the 2 later components increased monotonically with pulsing frequency (Figure 4). The magnitude of the responses was also modulated by the phase of the slow oscillation at which the stimulation train occurred: when VNS was delivered at the hyperpolarized phase (0-π rad) of the oscillation the VEPs were smaller than the ones elicited by the VNS occurring at the depolarized phase (π-2π rad) (Figure 4B). These results suggested that VNS during NREM sleep evokes dose- and phase-dependent, SO-like responses in the cortex.

Importantly, the number of transitions between brain states was not altered by the presence of stimulation https://doi.org/10.6084/m9.figshare.12724739.v6), at least with the relatively short VNS train tested in our study. Because our brain state classification algorithm was based on the power of different frequency bands, this result indicated that VNS did not affect the power in any frequency band as well as it did not affect coherence (Figure 7, S10, https://doi.org/10.6084/m9.figshare.12724739.v6). Among other sleep parameters, changes in the duration of REM and slow-wave sleep cycles have been described in association with clinical VNS, typically delivered in 30 s-ON/5 min-OFF periods (Romero-Osorioet al., 2018). Our results suggest that shorter ON periods may be less prone to affecting sleep cycles.

In contrast to the 2 later components, the early VEP component did not change between awake and sleep states (Figure 4). This difference could be explained by activation of separate circuits by VNS. The early VEP component is likely initiated by direct activation of large, somatic afferents or it could be myogenic in origin (Hammond EJ *et al*. 1992). It could represent the activation of a relatively direct anatomical pathway involving the nucleus of the solitary tract and possibly other first or second-order nuclei that project to the thalamus and cortex (Berthoud HR and WL Neuhuber 2000; Gamboa-Esteves FO et al. 2001). In addition to motor innervation of laryngeal and pharyngeal muscles, the vagus also provides afferent innervation to other structures of the larynx, including somatosensory innervation of laryngeal and pharyngeal mucosa and proprioceptive innervation of laryngeal muscles and joints (Puizillout J-J 2005). Stimulus-elicited contraction of laryngeal muscles could activate those afferents, thereby producing afferent volleys that manifest as stimulus-evoked potentials in sensorimotor cortical areas (Sasaki R et al. 2017). This would explain the disappearance of VEPs when muscle contraction was blocked (Hammond EJ *et al*. 1992).

The later VEP components, on the other hand, could reflect activation of longer polysynaptic pathways, mediated by relays in the brainstem, midbrain, hypothalamus, hippocampus, thalamus and cortex (Henry TR 2002). In contrast to the early component, later components of the VEPs showed dose-dependence, suggesting more significant temporal synaptic summation, consistent with a polysynaptic pathway. Although the cervical vagus involves sensory fiber populations with different sizes, myelination properties and conduction velocities (Agostoni E et al. 1957), it is unlikely that activation of faster or slower fibers alone can account for the different VEP components, since the different conduction velocities give rise to latency differences that are at least one order of magnitude smaller than the latencies seen in the VEPs. Even though there were instances of polarity reversal across brain states, those were limited to the intermediate-latency VEP component; in those cases, those components were relatively small in amplitude (<50uV peak amplitude) (e.g. Figure 5A, middle panel, for both subjects). This likely reflects the heterogeneity and variability of the waves included in the intermediate-latency component and the fact that only one of them (the largest in each VEP) contributed to the reported measurement: sometimes a positive wave was the largest, whereas in other instances the negative was the largest.

### Possible mechanisms mediating brain-state modulation of VEPs

Brainstem and midbrain areas that receive afferent inputs from the NTS largely project to the cortex via the thalamus (Berthoud HR and WL Neuhuber 2000; Henry TR 2002). Therefore the large-scale changes in thalamo-cortical circuits occurring during sleep (McCormick DA and T Bal 1997) could play a role in modulating the cortical responses elicited by VNS. The thalamus is a major gateway into the cerebral cortex and the first station at which incoming signals can be blocked by synaptic inhibition during sleep. Thalamo-cortical and cortico-cortical interactions contribute to the changes that brain activity undergoes during the switch from an aroused state, more receptive to “external” signals, to a more isolated sleep state, which is driven by “internal”, oscillatory activity (McCormick DA and T Bal 1997; Sanchez-Vives MV and DA McCormick 2000; Steriade M 2004). The brain state dependence of VEPs suggests that the effect of ascending volleys generated by VNS on cortical activity is shaped by the state of ongoing thalamo-cortical and cortico-cortical interactions, much like other sensory evoked potentials. During NREM sleep K-complexes and vertex sharp waves can be evoked by auditory or other sensory stimuli (Colrain IM et al. 1999; Colrain IM, P Di Parsia, et al. 2000; Colrain IM, KE Webster, et al. 2000). Likewise, cortical TMS pulses delivered during sleep trigger the generation of delta waves (Massimini M *et al*. 2007). Therefore, during sleep, slow oscillations, K-complexes and vertex sharp waves, could all contribute to the responses evoked by stimulation, manifesting as larger intermediate- and long-latency VEPs. The fact that the relatively long latency, slower components of the VEPs were the ones mostly augmented during NREM sleep, agrees with the shift to slower spontaneous EEG components in that sleep stage, such as delta waves and K-complexes. The shift to larger in amplitude and slower in time-course stimulus-evoked and spontaneous signatures of cortical activity during NREM may reflect neuronal synchronization across larger cortical and subcortical neuronal populations, which has been demonstrated in that sleep stage (Scammell TE et al. 2017).

Interestingly, several studies demonstrated that the balance between parasympathetic and sympathetic activity changes during sleep. Spectral analysis of heart rate variability, a measure of autonomic activity, showed an increase of parasympathetic tone during NREM sleep (Berlad II et al. 1993; Trinder J et al. 2001; Mendez M et al. 2006; Cabiddu R et al. 2012). Increased vagal tone could be mediated by increased efferent vagal activity, but also by increased responsiveness of the afferent vagus to peripheral stimuli (Laborde S et al. 2018). Thus, increased vagal tone during sleep might contribute to a larger VEP compared to evoked responses elicited by the same stimuli during waking. Although these studies indicate that the autonomic system in specifically affected by changes in behavioral states, our control experiments on a third monkey suggested that the brain state modulation of cortical evoked potentials is not only specific to the stimulation of the vagus, but it is visible also when the stimulation is delivered to another peripheral nerve (Figure S7, https://doi.org/10.6084/m9.figshare.12724739.v6), or in humans when it is delivered to the cortex through TMS or intracortical stimulation (Massimini M *et al*. 2007; Pigorini A et al. 2015). Because the non-specificity to the kind of stimulation it is possible that this phenomenon is related to the characteristic bistability of the corticothalamic network during sleep (Sanchez-Vives MV and DA McCormick 2000), which is amplified by a strong initial activation. At the cellular level the bistability of NREM sleep is thought to be primarily due to the dynamics of potassium current (K+), which could lead neurons into a hyperpolarized, silent state followed a strong activation (Compte A et al. 2003), which based on our results could also be triggered by stimulating peripheral nerves.

Recently, it was shown that VNS in mice engages cholinergic and adrenergic axons to the cortex, resulting in wide-spread activation of excitatory cortical neurons and changes in the state of arousal (Collins L *et al*. 2021). Importantly, in that study, VNS-triggered cortical activation persisted even in light anesthesia, in the absence of motor activity, indicating that part of the cortical effects of VNS are not brain-state dependent. Part of those non-state-dependent cortical effects may correspond to the earlier components of the VEP responses we documented in our study. It is likely that ascending volleys triggered by VNS follow both oligosynaptic and multisynaptic pathways, result in a mix of fast and slow synaptic processes in different cell types and interact differently with modes of ongoing cortical activity. Ultimately, more hypothesis-driven experiments could shed light into the mechanisms of brain-state modulation of cortical actions of VNS. Local application of agonists or antagonists of cholinergic, noradrenergic and glutamatergic neurotransmission in brainstem, midbrain or thalamocortical sites, in conjunction with brain recordings and behavioral and anesthesia state manipulations can help resolve the relative contribution of these chemical systems and brain state to cortical responses to VNS. For example, if local cortical application of antagonists of cholinergic neurotransmission does not result in changes in VEP amplitudes, that would suggest that VEPs reflect cortical activation by mostly glutamatergic thalamocortical projections. Alternatively, cholinergic antagonists could reduce the amplitude of early components of VEPs, which are largely unaffected by brain state, or of late components of VEPs, which are heavily modulated by brain state. Such experiments can dissociate the contribution to the shape and the state modulation of VEPs of ascending stimulus-triggered volleys and of ongoing cortical dynamics. In rodent models of VNS, there are additional experimental options, including wide field optical imaging of large cortical and subcortical neuronal populations in response to VNS (e.g. (Collins L *et al*. 2021)) and localized optogenetic activation (or block) of such systems instead of (or concurrently with) electrical VNS.

### Effect of pulsing frequency on cortical responses to VNS

In this study we delivered trains of pulses at pulsing frequencies ranging from 5 Hz to 300 Hz. For all brain states, VEPs recorded from different cortical areas had higher amplitudes at the highest pulsing frequency (300 Hz) than at the lowest pulsing frequency (5 Hz) (Figure 3). Monkey M1, in particular, showed a monotonic increase in the magnitude of the 2 later components with increasing pulsing frequency (Figure 4). Such monotonic relationship is different from the inverted-U-shaped relationship to a number of brain function readouts described previously. For example, a pulsing frequency around 30Hz resulted in an increased cortical map plasticity, whereas higher or lower VNS frequencies failed to induce plasticity effects (Buell EP et al. 2018; Buell EP et al. 2019). Initial clinical studies of VNS used pulsing frequencies of 30 or 50 Hz (Uthman BM et al. 1993), and in subsequent pivotal trials and in current clinical recommendations the pulsing frequency has been typically 20 or 30 Hz (<u>Cyberonics 2013, November 17 VNS Therapy Products</u>. Retrieved from VNS Therapy System Physicians’s Manual (US): http://dynamic.cyberonics.com/manuals/).

The reasons for this discrepancy are not clear. It could be due to the differences in number of VNS pulses delivered, but it could also be that the read-out of VNS in our study, evoked cortical activity, correlates poorly with VNS outcomes used in other studies, like cortical map plasticity, behavioral recovery or long-term suppression of epileptic activity. Our results could in principle be explained by temporal summation of synaptic responses to VNS. For example, temporal summation could happen at the level of NTS; in monosynaptically-driven NTS responses, excitatory post-synaptic potentials last for 10-20 ms, and therefore temporal summation would occur at frequencies above 50-100 Hz (Austgen JR et al. 2011). A monotonic relationship between VNS pulsing frequency and neuronal firing in locus coeruleus has been described (Hulsey DR et al. 2017), suggesting a temporal summation mechanism for frequency dependency. This interpretation is also supported by our control experiment on a third animal, in which the median nerve was stimulated: the elicited evoked responses by single pulses were smaller and less modulated by brain state than the responses elicited by 5-pulse trains at 300Hz (Figure S7, https://doi.org/10.6084/m9.figshare.12724739.v6).

However, simple temporal summation is probably not the only mechanism behind frequency dependency of VEPs. Central cholinergic and noradrenergic pathways engage a variety of cell types in the cortex, thalamus and subcortical areas (e.g. (Picciotto MR et al. 2012)) and the net effect on cell excitability depends on the receptor subtypes, their location on the cells and on the mode of signaling (e.g. traditional synaptic signaling occurring locally vs. diffusion-based volume transmission occurring at a distance). Given the breadth and heterogeneity of brain networks activated by VNS, it is likely that the frequency dependency arises from interactions between synaptic temporal summation, the activation of fast, excitatory and slower, inhibitory ionotropic receptors, the fast time course of synaptic signaling vs. the slower, distant volume effects of neurotransmitter release, the conduction delays of central pathways and thalamocortical loops engaged by VNS and the properties of ongoing firing of engaged neurons at different cortical areas.

### VNS and targeted neuroplasticity

Studies in animal models have shown that electrically stimulating the vagus nerve leads to a release of plasticity-related neuromodulators in the brain, including acetylcholine and norepinephrine (Nichols JA *et al*. 2011). Those neuromodulators regulate plasticity by acting as ‘on-off switches’ which enable plastic changes to occur by engaging synaptic processes (Kilgard MP and MM Merzenich 1998; Sara SJ 2009; Sara SJ and S Bouret 2012). Sleep has been shown to play a crucial role for skill learning and memory consolidation (Huber R et al. 2004; Rasch B and J Born 2013; Gulati T et al. 2014) and direct manipulation of brain activity during sleep can result in changes in task performance (Gulati T et al. 2017; Rembado I et al. 2017; Ketz N et al. 2018; Kim J et al. 2019). Although the mechanisms underlying off-line learning are not entirely understood, one possibility involves the autonomic nervous system (ANS). In particular, Whitehurst et al. (Whitehurst LN et al. 2016) showed that improvements in tests of associative memory were associated with vagal-mediated ANS activity during REM sleep. Stimulation of the vagus nerve affects task performance when it is paired with active training (Pruitt DT *et al*. 2016). It is unknown whether VNS could have different cognitive effects when delivered during different brain states, including sleep. Our findings argue that this could be a possibility. If true, this would have implications in the use of VNS, and other methods of neuromodulation, to enhance neuroplasticity in healthy subjects and in patients with neurological disease.

### VNS and interoception

The vagus is the main conduit for interoceptive sensory signals, supporting conscious and unconscious perception of bodily events and states (Paciorek A and L Skora 2020). Afferent visceral signals related to physiological events, like heart rhythm and breathing, are conveyed by the vagus and elicit event-related potentials that are measurable with intracranial EEG. Examples are the heartbeat-evoked cortical potential (Park HD et al. 2018) and cortical activity related to breathing (Herrero JL et al. 2018). Such cortical signatures of visceral physiology reflect population-level aspects of processes by which the brain integrates and interprets interoceptive information (Paciorek A and L Skora 2020). It is unknown how these signatures are modulated by ongoing cortical activity and our study offers some insight. The fact that the same vagal afferent volley leads to different cortical responses depending on brain state (e.g. smaller response during wakefulness than during NREM sleep) indicates that vagal interoceptive information conveyed at different times of day and during different mental and behavioral states may shape how the continuous stream of visceral signals affects motivational states, adaptive behavior and emotion (Critchley HD and NA Harrison 2013).

### Implications for VNS therapies

Our study has implications for current and future VNS therapies. It is unknown whether there is a relationship between the size and amplitude of VEPs and the anti-seizure effect of VNS. However, to the extent that, for a given brain state, larger VEPs represent stronger stimulus-elicited activation of cortical circuits, it is reasonable to consider VEPs as markers of the degree of engagement by stimuli of fibers in the vagus and vagal pathways in the brain. In that sense, VEPs recorded via scalp EEG could be used clinically as a physiological marker to estimate nerve fiber engagement, especially of large vagal afferents, for which there are no good marker candidates, in contrast to small vagal afferents that can be assessed by stimulus-elicited changes in breathing (Chang YC et al. 2020). Resolving the engagement of large vagal afferents in real time would be key to the clinical calibration of VNS therapies targeting brain diseases in individual patients. Second, to the extent that modulation of VEPs by brain state represents changes in the excitability of the ascending vagal pathways and/or the cortical populations receiving ascending stimulus volleys, our findings may be relevant to the optimization of VNS therapies with regard to time of day or brain state, in the context of closed-loop neuromodulation (Zanos S 2019). Finally, our finding that short VNS trains with higher pulsing frequencies, especially above 30-50 Hz, result in larger VEPs indicates that this high-frequency “burst-mode” VNS can exert robust, transient effects on cortical activity that could be useful in applications of VNS in which neuromodulation has to be delivered with high temporal accuracy. Such short trains may also lead to fewer off-target effects from peripheral organs, such as from the heart (Chang YC *et al*. 2020).

## Supporting information

https://doi.org/10.6084/m9.figshare.12724739.v6

## Acknowledgments

This work was supported by NIH grants RO1 NS12542 and RR00166 to E.F.

## Author contributions

I.R. collected data, analyzed data, interpreted results, wrote the paper

W.S. analyzed data

D.S. performed surgeries for vagus nerve implant

A.L. collected data

L.S. hardware and software development of Neurochip

E.F., S.P. interpreted results

S.Z. performed surgeries for cortical implant, designed experiments, interpreted results, wrote the paper

## References

Agostoni E, Chinnock JE, De Daly MB, Murray JG. 1957. Functional and histological studies of the vagus nerve and its branches to the heart, lungs and abdominal viscera in the cat. J Physiol. 135:182–205.

Armitage R. 1995. The distribution of EEG frequencies in REM and NREM sleep stages in healthy young adults. Sleep. 18:334–341.

Austgen JR, Hermann GE, Dantzler HA, Rogers RC, Kline DD. 2011. Hydrogen sulfide augments synaptic neurotransmission in the nucleus of the solitary tract. J Neurophysiol. 106:1822–1832.

Babadi B, Brown EN. 2014. A review of multitaper spectral analysis. IEEE Trans Biomed Eng. 61:1555–1564.

Berlad II, Shlitner A, Ben-Haim S, Lavie P. 1993. Power spectrum analysis and heart rate variability in Stage 4 and REM sleep: evidence for state-specific changes in autonomic dominance. J Sleep Res. 2:88–90.

Berthoud HR, Neuhuber WL. 2000. Functional and chemical anatomy of the afferent vagal system. Auton Neurosci. 85:1–17.

Buell EP, Borland MS, Loerwald KW, Chandler C, Hays SA, Engineer CT, Kilgard MP. 2019. Vagus Nerve Stimulation Rate and Duration Determine whether Sensory Pairing Produces Neural Plasticity. Neuroscience. 406:290–299.

Buell EP, Loerwald KW, Engineer CT, Borland MS, Buell JM, Kelly CA, Khan, II, Hays SA, Kilgard MP. 2018. Cortical map plasticity as a function of vagus nerve stimulation rate. Brain Stimul. 11:1218–1224.

Cabiddu R, Cerutti S, Viardot G, Werner S, Bianchi AM. 2012. Modulation of the Sympatho-Vagal Balance during Sleep: Frequency Domain Study of Heart Rate Variability and Respiration. Front Physiol. 3:45.

Cao B, Wang J, Shahed M, Jelfs B, Chan RH, Li Y. 2016. Vagus Nerve Stimulation Alters Phase Synchrony of the Anterior Cingulate Cortex and Facilitates Decision Making in Rats. Sci Rep. 6:35135.

Car A, Jean A, Roman C. 1975. A pontine primary relay for ascending projections of the superior laryngeal nerve. Exp Brain Res. 22:197–210.

Chang YC, Cracchiolo M, Ahmed U, Mughrabi I, Gabalski A, Daytz A, Rieth L, Becker L, Datta-Chaudhuri T, Al-Abed Y, Zanos TP, Zanos S. 2020. Quantitative estimation of nerve fiber engagement by vagus nerve stimulation using physiological markers. Brain Stimul. 13:1617–1630.

Charbonnier S, Zoubek L, Lesecq S, Chapotot F. 2011. Self-evaluated automatic classifier as a decision-support tool for sleep/wake staging. Comput Biol Med. 41:380–389.

Cheyuo C, Jacob A, Wu R, Zhou M, Coppa GF, Wang P. 2011. The parasympathetic nervous system in the quest for stroke therapeutics. J Cereb Blood Flow Metab. 31:1187–1195.

Collins L, Boddington L, Steffan PJ, McCormick D. 2021. Vagus nerve stimulation induces widespread cortical and behavioral activation. Curr Biol.

Colrain IM, Di Parsia P, Gora J. 2000. The impact of prestimulus EEG frequency on auditory evoked potentials during sleep onset. Can J Exp Psychol. 54:243–254.

Colrain IM, Webster KE, Hirst G. 1999. The N550 component of the evoked K-complex: a modality non-specific response? J Sleep Res. 8:273–280.

Colrain IM, Webster KE, Hirst G, Campbell KB. 2000. The roles of vertex sharp waves and K-complexes in the generation of N300 in auditory and respiratory-related evoked potentials during early stage 2 NREM sleep. Sleep. 23:97–106.

Compte A, Sanchez-Vives MV, McCormick DA, Wang XJ. 2003. Cellular and network mechanisms of slow oscillatory activity (<1 Hz) and wave propagations in a cortical network model. J Neurophysiol. 89:2707–2725.

Critchley HD, Garfinkel SN. 2017. Interoception and emotion. Curr Opin Psychol. 17:7–14.

Critchley HD, Harrison NA. 2013. Visceral influences on brain and behavior. Neuron. 77:624–638.

Daley JT, Turner RS, Freeman A, Bliwise DL, Rye DB. 2006. Prolonged assessment of sleep and daytime sleepiness in unrestrained Macaca mulatta. Sleep. 29:221–231.

Dawson J, Pierce D, Dixit A, Kimberley TJ, Robertson M, Tarver B, Hilmi O, McLean J, Forbes K, Kilgard MP, Rennaker RL, Cramer SC, Walters M, Engineer N. 2016. Safety, Feasibility, and Efficacy of Vagus Nerve Stimulation Paired With Upper-Limb Rehabilitation After Ischemic Stroke. Stroke. 47:143–150.

Destexhe A, Contreras D, Steriade M. 1999. Spatiotemporal analysis of local field potentials and unit discharges in cat cerebral cortex during natural wake and sleep states. J Neurosci. 19:4595–4608.

Di Lazzaro V, Oliviero A, Pilato F, Saturno E, Dileone M, Meglio M, Colicchio G, Barba C, Papacci F, Tonali PA. 2004. Effects of vagus nerve stimulation on cortical excitability in epileptic patients. Neurology. 62:2310–2312.

Elger G, Hoppe C, Falkai P, Rush AJ, Elger CE. 2000. Vagus nerve stimulation is associated with mood improvements in epilepsy patients. Epilepsy Res. 42:203–210.

Engineer CT, Engineer ND, Riley JR, Seale JD, Kilgard MP. 2015. Pairing Speech Sounds With Vagus Nerve Stimulation Drives Stimulus-specific Cortical Plasticity. Brain Stimul. 8:637–644.

Engineer ND, Riley JR, Seale JD, Vrana WA, Shetake JA, Sudanagunta SP, Borland MS, Kilgard MP. 2011. Reversing pathological neural activity using targeted plasticity. Nature. 470:101–104.

Fallgatter AJ, Ehlis AC, Ringel TM, Herrmann MJ. 2005. Age effect on far field potentials from the brain stem after transcutaneous vagus nerve stimulation. Int J Psychophysiol. 56:37–43.

Gamboa-Esteves FO, Lima D, Batten TF. 2001. Neurochemistry of superficial spinal neurones projecting to nucleus of the solitary tract that express c-fos on chemical somatic and visceral nociceptive input in the rat. Metab Brain Dis. 16:151–164.

Gasser T, Bacher P, Mocks J. 1982. Transformations towards the normal distribution of broad band spectral parameters of the EEG. Electroencephalogr Clin Neurophysiol. 53:119–124.

George MS, Nahas Z, Bohning DE, Kozel FA, Anderson B, Chae JH, Lomarev M, Denslow S, Li X, Mu C. 2002. Vagus nerve stimulation therapy: a research update. Neurology. 59:S56–61.

Gulati T, Guo L, Ramanathan DS, Bodepudi A, Ganguly K. 2017. Neural reactivations during sleep determine network credit assignment. Nat Neurosci. 20:1277–1284.

Gulati T, Ramanathan DS, Wong CC, Ganguly K. 2014. Reactivation of emergent task-related ensembles during slow-wave sleep after neuroprosthetic learning. Nat Neurosci. 17:1107–1113.

Hagen K, Ehlis AC, Schneider S, Haeussinger FB, Fallgatter AJ, Metzger FG. 2014. Influence of different stimulation parameters on the somatosensory evoked potentials of the nervus vagus--how varied stimulation parameters affect VSEP. J Clin Neurophysiol. 31:143–148.

Hammond EJ, Uthman BM, Reid SA, Wilder BJ. 1992. Electrophysiologic studies of cervical vagus nerve stimulation in humans: II. Evoked potentials. Epilepsia. 33:1021–1028.

Hammond EJ, Uthman BM, Reid SA, Wilder BJ. 1992. Electrophysiological studies of cervical vagus nerve stimulation in humans: I. EEG effects. Epilepsia. 33:1013–1020.

Hassert DL, Miyashita T, Williams CL. 2004. The effects of peripheral vagal nerve stimulation at a memory-modulating intensity on norepinephrine output in the basolateral amygdala. Behav Neurosci. 118:79–88.

Hays SA, Ruiz A, Bethea T, Khodaparast N, Carmel JB, Rennaker RL, 2nd, Kilgard MP. 2016. Vagus nerve stimulation during rehabilitative training enhances recovery of forelimb function after ischemic stroke in aged rats. Neurobiol Aging. 43:111–118.

Hennevin E, Huetz C, Edeline JM. 2007. Neural representations during sleep: from sensory processing to memory traces. Neurobiol Learn Mem. 87:416–440.

Henry TR. 2002. Therapeutic mechanisms of vagus nerve stimulation. Neurology. 59:S3–14.

Herrero JL, Khuvis S, Yeagle E, Cerf M, Mehta AD. 2018. Breathing above the brain stem: volitional control and attentional modulation in humans. J Neurophysiol. 119:145–159.

Hsieh KC, Robinson EL, Fuller CA. 2008. Sleep architecture in unrestrained rhesus monkeys (Macaca mulatta) synchronized to 24-hour light-dark cycles. Sleep. 31:1239–1250.

Huber R, Ghilardi MF, Massimini M, Tononi G. 2004. Local sleep and learning. Nature. 430:78–81.

Hulsey DR, Riley JR, Loerwald KW, Rennaker RL, 2nd, Kilgard MP, Hays SA. 2017. Parametric characterization of neural activity in the locus coeruleus in response to vagus nerve stimulation. Exp Neurol. 289:21–30.

Ito S, Craig AD. 2005. Vagal-evoked activity in the parafascicular nucleus of the primate thalamus. J Neurophysiol. 94:2976–2982.

Ketz N, Jones AP, Bryant NB, Clark VP, Pilly PK. 2018. Closed-Loop Slow-Wave tACS Improves Sleep-Dependent Long-Term Memory Generalization by Modulating Endogenous Oscillations. J Neurosci. 38:7314–7326.

Khalsa SS, Adolphs R, Cameron OG, Critchley HD, Davenport PW, Feinstein JS, Feusner JD, Garfinkel SN, Lane RD, Mehling WE, Meuret AE, Nemeroff CB, Oppenheimer S, Petzschner FH, Pollatos O, Rhudy JL, Schramm LP, Simmons WK, Stein MB, Stephan KE, Van den Bergh O, Van Diest I, von Leupoldt A, Paulus MP, Interoception Summit p. 2018. Interoception and Mental Health: A Roadmap. Biol Psychiatry Cogn Neurosci Neuroimaging. 3:501–513.

Kilgard MP, Merzenich MM. 1998. Cortical map reorganization enabled by nucleus basalis activity. Science. 279:1714–1718.

Kim J, Gulati T, Ganguly K. 2019. Competing Roles of Slow Oscillations and Delta Waves in Memory Consolidation versus Forgetting. Cell. 179:514–526 e513.

Krakovska A, Mezeiova K. 2011. Automatic sleep scoring: a search for an optimal combination of measures. Artif Intell Med. 53:25–33.

Laborde S, Mosley E, Mertgen A. 2018. Vagal Tank Theory: The Three Rs of Cardiac Vagal Control Functioning - Resting, Reactivity, and Recovery. Front Neurosci. 12:458.

Livingstone MS, Hubel DH. 1981. Effects of sleep and arousal on the processing of visual information in the cat. Nature. 291:554–561.

Marshall AC, Gentsch A, Schutz-Bosbach S. 2018. The Interaction between Interoceptive and Action States within a Framework of Predictive Coding. Front Psychol. 9:180.

Mascetti GG. 2016. Unihemispheric sleep and asymmetrical sleep: behavioral, neurophysiological, and functional perspectives. Nat Sci Sleep. 8:221–238.

Massimini M, Ferrarelli F, Esser SK, Riedner BA, Huber R, Murphy M, Peterson MJ, Tononi G. 2007. Triggering sleep slow waves by transcranial magnetic stimulation. Proc Natl Acad Sci U S A. 104:8496–8501.

Massimini M, Ferrarelli F, Huber R, Esser SK, Singh H, Tononi G. 2005. Breakdown of cortical effective connectivity during sleep. Science. 309:2228–2232.

McCormick DA, Bal T. 1997. Sleep and arousal: thalamocortical mechanisms. Annu Rev Neurosci. 20:185–215.

Mendez M, Bianchi AM, Villantieri O, Cerutti S. 2006. Time-varying analysis of the heart rate variability during REM and non REM sleep stages. Conf Proc IEEE Eng Med Biol Soc. 1:3576–3579.

Metzger FG, Polak T, Aghazadeh Y, Ehlis AC, Hagen K, Fallgatter AJ. 2012. Vagus somatosensory evoked potentials--a possibility for diagnostic improvement in patients with mild cognitive impairment? Dement Geriatr Cogn Disord. 33:289–296.

Nichols JA, Nichols AR, Smirnakis SM, Engineer ND, Kilgard MP, Atzori M. 2011. Vagus nerve stimulation modulates cortical synchrony and excitability through the activation of muscarinic receptors. Neuroscience. 189:207–214.

Noble LJ, Gonzalez IJ, Meruva VB, Callahan KA, Belfort BD, Ramanathan KR, Meyers E, Kilgard MP, Rennaker RL, McIntyre CK. 2017. Effects of vagus nerve stimulation on extinction of conditioned fear and post-traumatic stress disorder symptoms in rats. Transl Psychiatry. 7:e1217.

Paciorek A, Skora L. 2020. Vagus Nerve Stimulation as a Gateway to Interoception. Front Psychol. 11:1659.

Panagiotou M, Vyazovskiy VV, Meijer JH, Deboer T. 2017. Differences in electroencephalographic non-rapid-eye movement sleep slow-wave characteristics between young and old mice. Sci Rep. 7:43656.

Park HD, Bernasconi F, Salomon R, Tallon-Baudry C, Spinelli L, Seeck M, Schaller K, Blanke O. 2018. Neural Sources and Underlying Mechanisms of Neural Responses to Heartbeats, and their Role in Bodily Self-consciousness: An Intracranial EEG Study. Cereb Cortex. 28:2351–2364.

Pfurtscheller G, Stancak A, Jr., Neuper C. 1996. Event-related synchronization (ERS) in the alpha band--an electrophysiological correlate of cortical idling: a review. Int J Psychophysiol. 24:39–46.

Picciotto MR, Higley MJ, Mineur YS. 2012. Acetylcholine as a neuromodulator: cholinergic signaling shapes nervous system function and behavior. Neuron. 76:116–129.

Pigorini A, Sarasso S, Proserpio P, Szymanski C, Arnulfo G, Casarotto S, Fecchio M, Rosanova M, Mariotti M, Lo Russo G, Palva JM, Nobili L, Massimini M. 2015. Bistability breaks-off deterministic responses to intracortical stimulation during non-REM sleep. Neuroimage. 112:105–113.

Polak T, Ehlis AC, Langer JB, Plichta MM, Metzger F, Ringel TM, Fallgatter AJ. 2007. Non-invasive measurement of vagus activity in the brainstem - a methodological progress towards earlier diagnosis of dementias? J Neural Transm (Vienna). 114:613–619.

Polak T, Herrmann MJ, Muller LD, Zeller JBM, Katzorke A, Fischer M, Spielmann F, Weinmann E, Hommers L, Lauer M, Fallgatter AJ, Deckert J. 2017. Near-infrared spectroscopy (NIRS) and vagus somatosensory evoked potentials (VSEP) in the early diagnosis of Alzheimer’s disease: rationale, design, methods, and first baseline data of the Vogel study. J Neural Transm (Vienna). 124:1473–1488.

Polak T, Markulin F, Ehlis AC, Langer JB, Ringel TM, Fallgatter AJ. 2009. Far field potentials from brain stem after transcutaneous vagus nerve stimulation: optimization of stimulation and recording parameters. J Neural Transm (Vienna). 116:1237–1242.

Porter BA, Khodaparast N, Fayyaz T, Cheung RJ, Ahmed SS, Vrana WA, Rennaker RL, 2nd, Kilgard MP. 2012. Repeatedly pairing vagus nerve stimulation with a movement reorganizes primary motor cortex. Cereb Cortex. 22:2365–2374.

Pritchard TC, Hamilton RB, Norgren R. 2000. Projections of the parabrachial nucleus in the old world monkey. Exp Neurol. 165:101–117.

Pruitt DT, Schmid AN, Kim LJ, Abe CM, Trieu JL, Choua C, Hays SA, Kilgard MP, Rennaker RL. 2016. Vagus Nerve Stimulation Delivered with Motor Training Enhances Recovery of Function after Traumatic Brain Injury. J Neurotrauma. 33:871–879.

Puizillout J-J. 2005. Central projections of vagal afferents: Publibook.

Qassim YT, Cutmore TR, James DA, Rowlands DD. 2013. Wavelet coherence of EEG signals for a visual oddball task. Comput Biol Med. 43:23–31.

Rachalski A, Authier S, Bassett L, Pouliot M, Tremblay G, Mongrain V. 2014. Sleep electroencephalographic characteristics of the Cynomolgus monkey measured by telemetry. J Sleep Res. 23:619–627.

Rasch B, Born J. 2013. About sleep’s role in memory. Physiol Rev. 93:681–766.

Ravan M, Sabesan S, D’Cruz O. 2017. On Quantitative Biomarkers of VNS Therapy Using EEG and ECG Signals. IEEE Trans Biomed Eng. 64:419–428.

Rembado I, Zanos S, Fetz EE. 2017. Cycle-Triggered Cortical Stimulation during Slow Wave Sleep Facilitates Learning a BMI Task: A Case Report in a Non-Human Primate. Front Behav Neurosci. 11:59.

Romcy-Pereira RN, de Araujo DB, Leite JP, Garcia-Cairasco N. 2008. A semi-automated algorithm for studying neuronal oscillatory patterns: a wavelet-based time frequency and coherence analysis. J Neurosci Methods. 167:384–392.

Sanchez-Vives MV, McCormick DA. 2000. Cellular and network mechanisms of rhythmic recurrent activity in neocortex. Nat Neurosci. 3:1027–1034.

Sara SJ. 2009. The locus coeruleus and noradrenergic modulation of cognition. Nat Rev Neurosci. 10:211–223.

Sara SJ, Bouret S. 2012. Orienting and reorienting: the locus coeruleus mediates cognition through arousal. Neuron. 76:130–141.

Sasaki R, Kotan S, Nakagawa M, Miyaguchi S, Kojima S, Saito K, Inukai Y, Onishi H. 2017. Presence and Absence of Muscle Contraction Elicited by Peripheral Nerve Electrical Stimulation Differentially Modulate Primary Motor Cortex Excitability. Front Hum Neurosci. 11:146.

Scammell TE, Arrigoni E, Lipton JO. 2017. Neural Circuitry of Wakefulness and Sleep. Neuron. 93:747–765.

Seth AK. 2013. Interoceptive inference, emotion, and the embodied self. Trends Cogn Sci. 17:565–573.

Shaw FZ, Lee SY, Chiu TH. 2006. Modulation of somatosensory evoked potentials during wake-sleep states and spike-wave discharges in the rat. Sleep. 29:285–293.

Shetake JA, Engineer ND, Vrana WA, Wolf JT, Kilgard MP. 2012. Pairing tone trains with vagus nerve stimulation induces temporal plasticity in auditory cortex. Exp Neurol. 233:342–349.

Sjogren MJ, Hellstrom PT, Jonsson MA, Runnerstam M, Silander HC, Ben-Menachem E. 2002. Cognition-enhancing effect of vagus nerve stimulation in patients with Alzheimer’s disease: a pilot study. J Clin Psychiatry. 63:972–980.

Steriade M. 1997. Synchronized activities of coupled oscillators in the cerebral cortex and thalamus at different levels of vigilance. Cereb Cortex. 7:583–604.

Steriade M. 2004. Acetylcholine systems and rhythmic activities during the waking--sleep cycle. Prog Brain Res. 145:179–196.

Steriade M, Nunez A, Amzica F. 1993. Intracellular analysis of relations between the slow (< 1 Hz) neocortical oscillation and other sleep rhythms of the electroencephalogram. J Neurosci. 13:3266–3283.

Steriade M, Nunez A, Amzica F. 1993. A novel slow (< 1 Hz) oscillation of neocortical neurons in vivo: depolarizing and hyperpolarizing components. J Neurosci. 13:3252–3265.

Trinder J, Kleiman J, Carrington M, Smith S, Breen S, Tan N, Kim Y. 2001. Autonomic activity during human sleep as a function of time and sleep stage. J Sleep Res. 10:253–264.

Tsakiris M, Critchley H. 2016. Interoception beyond homeostasis: affect, cognition and mental health. Philos Trans R Soc Lond B Biol Sci. 371.

Tyler R, Cacace A, Stocking C, Tarver B, Engineer N, Martin J, Deshpande A, Stecker N, Pereira M, Kilgard M, Burress C, Pierce D, Rennaker R, Vanneste S. 2017. Vagus Nerve Stimulation Paired with Tones for the Treatment of Tinnitus: A Prospective Randomized Double-blind Controlled Pilot Study in Humans. Sci Rep. 7:11960.

Upton AR, Tougas G, Talalla A, White A, Hudoba P, Fitzpatrick D, Clarke B, Hunt R. 1991. Neurophysiological effects of left vagal stimulation in man. Pacing Clin Electrophysiol. 14:70–76.

Uthman BM, Wilder BJ, Penry JK, Dean C, Ramsay RE, Reid SA, Hammond EJ, Tarver WB, Wernicke JF. 1993. Treatment of epilepsy by stimulation of the vagus nerve. Neurology. 43:1338–1345.

Vesuna S, Kauvar IV, Richman E, Gore F, Oskotsky T, Sava-Segal C, Luo L, Malenka RC, Henderson JM, Nuyujukian P, Parvizi J, Deisseroth K. 2020. Deep posteromedial cortical rhythm in dissociation. Nature. 586:87–94.

Vyazovskiy VV, Olcese U, Hanlon EC, Nir Y, Cirelli C, Tononi G. 2011. Local sleep in awake rats. Nature. 472:443–447.

Whitehurst LN, Cellini N, McDevitt EA, Duggan KA, Mednick SC. 2016. Autonomic activity during sleep predicts memory consolidation in humans. Proc Natl Acad Sci U S A. 113:7272–7277.

Zagon A, Kemeny AA. 2000. Slow hyperpolarization in cortical neurons: a possible mechanism behind vagus nerve simulation therapy for refractory epilepsy? Epilepsia. 41:1382–1389.

Zanos S. 2019. Closed-Loop Neuromodulation in Physiological and Translational Research. Cold Spring Harb Perspect Med. 9.

Zanos S, Moorjani S., Sabesan S., Fetz EE editor. Effects of vagus nerve stimulation on cortical activity and excitability in the nonhuman primate, SfN Annual Meeting, San Diego CA; 2016.

Zanos S, Richardson AG, Shupe L, Miles FP, Fetz EE. 2011. The Neurochip-2: an autonomous head-fixed computer for recording and stimulating in freely behaving monkeys. IEEE Trans Neural Syst Rehabil Eng. 19:427–435.

Zoubek L. CS, Lesecq S., Buguet A., Chapotot F. 2007. Feature selection for sleep/wake stages classification using data driven methods. Biomedical Signal Processing and Control. 2:171–179.

