## Supplementary material for "Cortical responses to vagus nerve stimulation are modulated by brain state in non-human primates": https://doi.org/10.6084/m9.figshare.12724739.v6

**Supplementary figures**

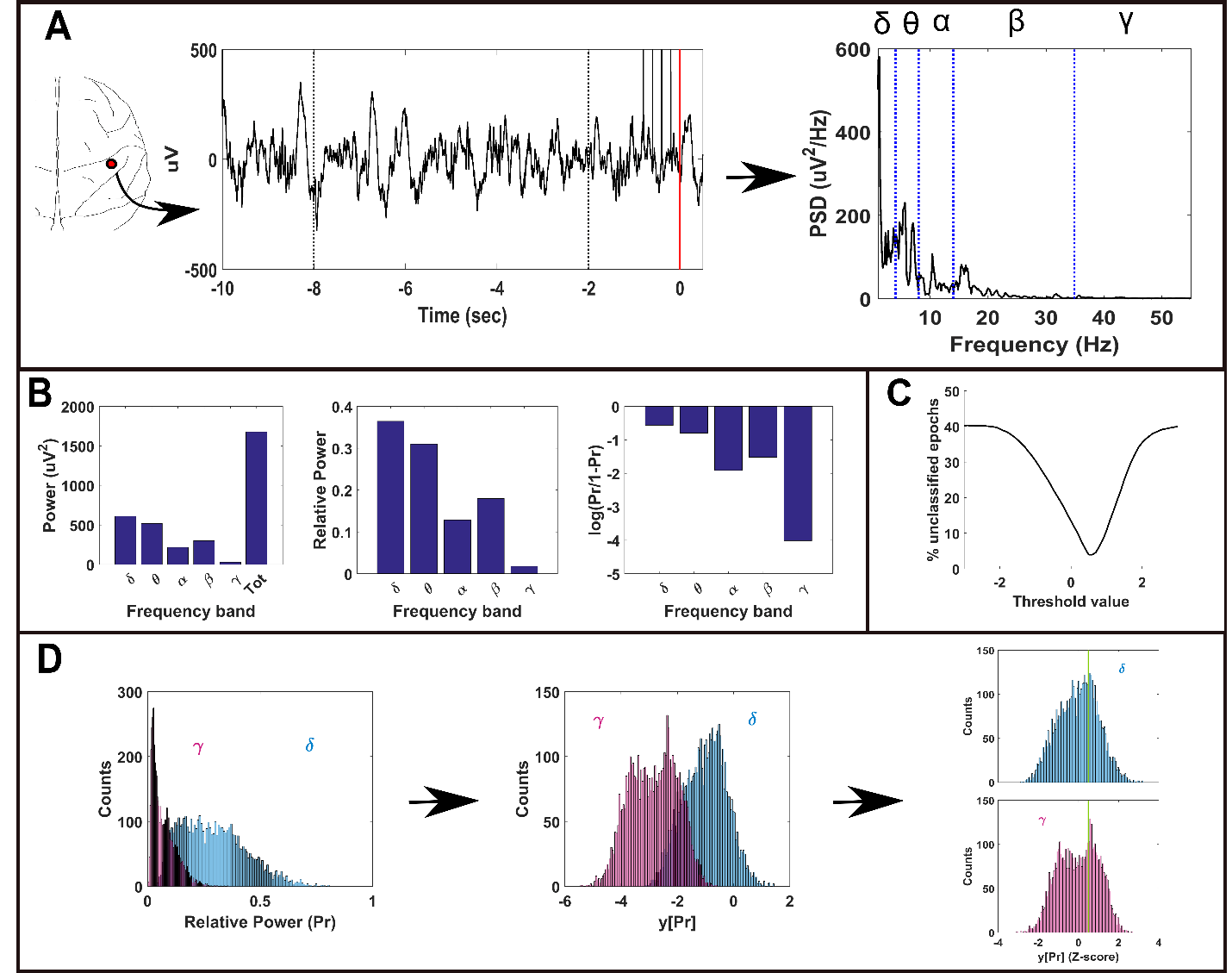

S1: Epoch based processing of ECoG power for classification of brain states. A) 10 seconds of raw signal before VNS of one cortical site (on the left). The red vertical line indicates the end of stimulation (the black vertical lines are the stimulation artifacts). The two vertical dashed lines mark the 6-sec epoch used to generate the power spectrum density on the right. The blue dashed lines mark the limits of the frequency bands considered for the power analysis (δ = 1-4 Hz, θ = 4-8 Hz, α = 8-14, β = 14-35 Hz, γ = 35-55 Hz). B) Example of power values for each frequency band calculated from the epoch displayed in A. From left to right: absolute power, relative power and transformed relative power. C) Number of unclassified epochs as a function of different power threshold values used to discriminate between brain states. The threshold which returned the minimum number of unclassified epochs was chosen for the classification procedure. D) Example of power distributions in delta and gamma bands generated by all epochs from one representative cortical site from one recording for three consecutive steps of the power processing. From left to right: distribution of relative power, transformed relative power and Z-scored transformed relative power (see Methods). The green vertical line in the last subplot points to the threshold value used to classify the brain states (i.e. minimum value of the curve shown in C).

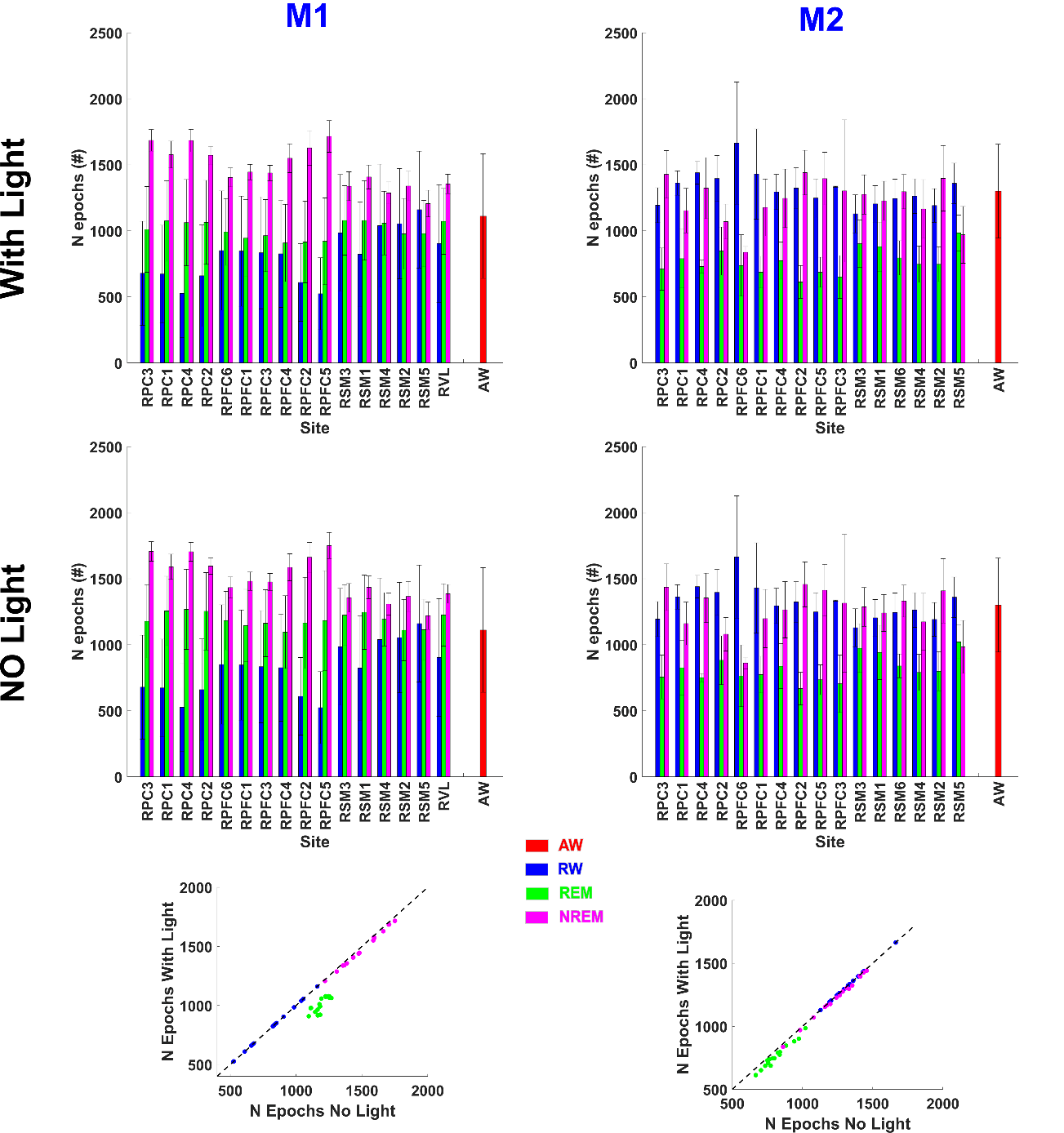

S2: Comparison based on the number of epochs for each classified brain state and for each recording site between two different classifiers. The first classifier included the information about when the room light was turned ON and OFF (top row) while the second classifier did not include this criterion (second row). The scatter plots on the bottom compare for each channel (represented by a dot) the number of classified epochs between the two classifiers. Despite some minor differences the two classifiers returned similar number of epochs for all channels, as highlighted by the identity line. Red: active-wake; blue: resting-wake; green: REM, pink: NREM.

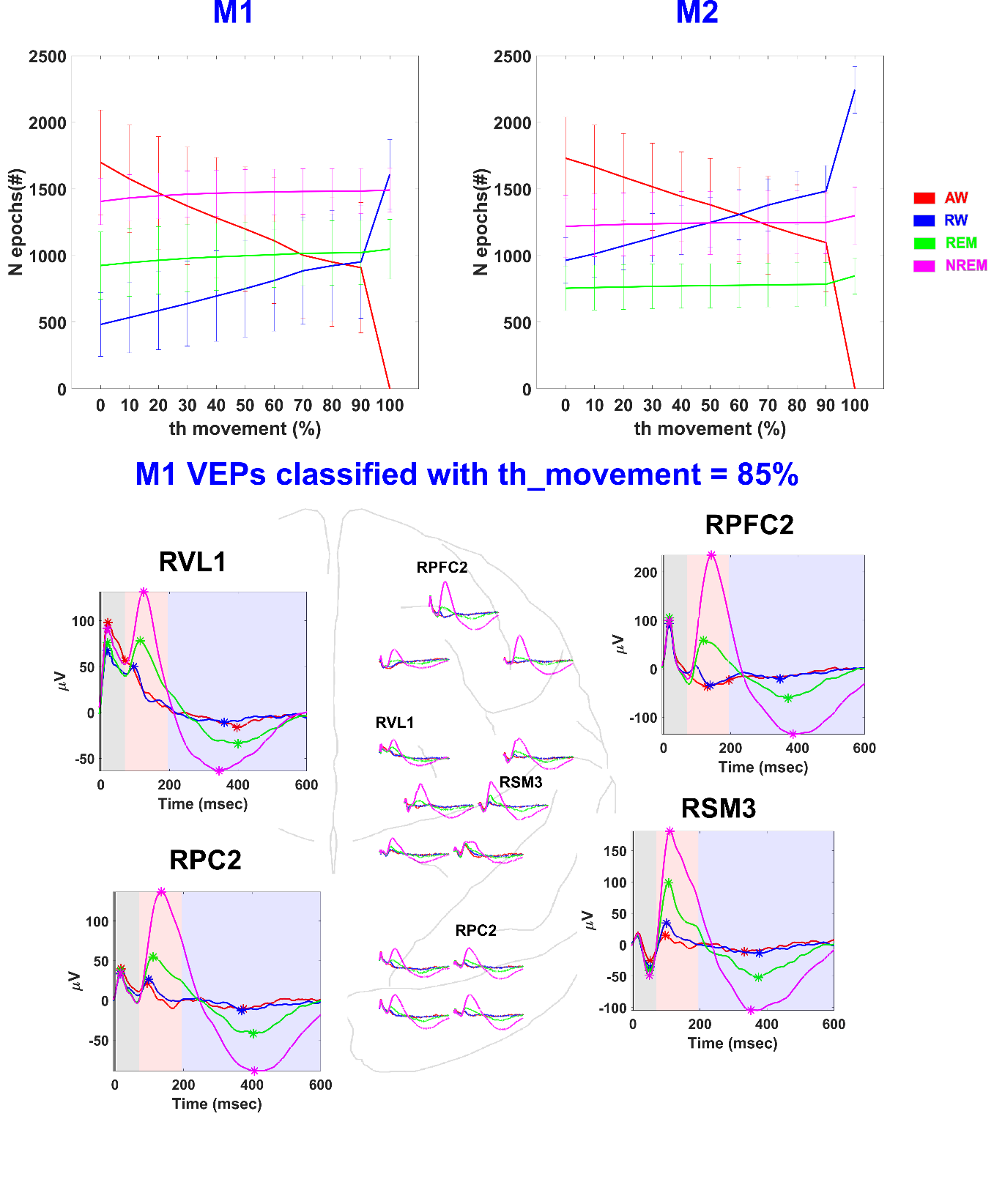

S3: Averaged number of classified epochs over all channels (± SD) as function of different movement threshold (X axis). We chose 60% for both animals to classify movement vs. resting. At 60% the number of epochs of classified as AW and RW are the same for M2. For M1 the two lines crossed at 85%. The morphology of the classified VEPs was not affected by the chosen movement threshold as shown in the bottom panel (compare it with the VEPs classified with Th_movement=60% shown in Figure 4).The number of classified epochs for REM and NREM is not significantly affected by the movement threshold.

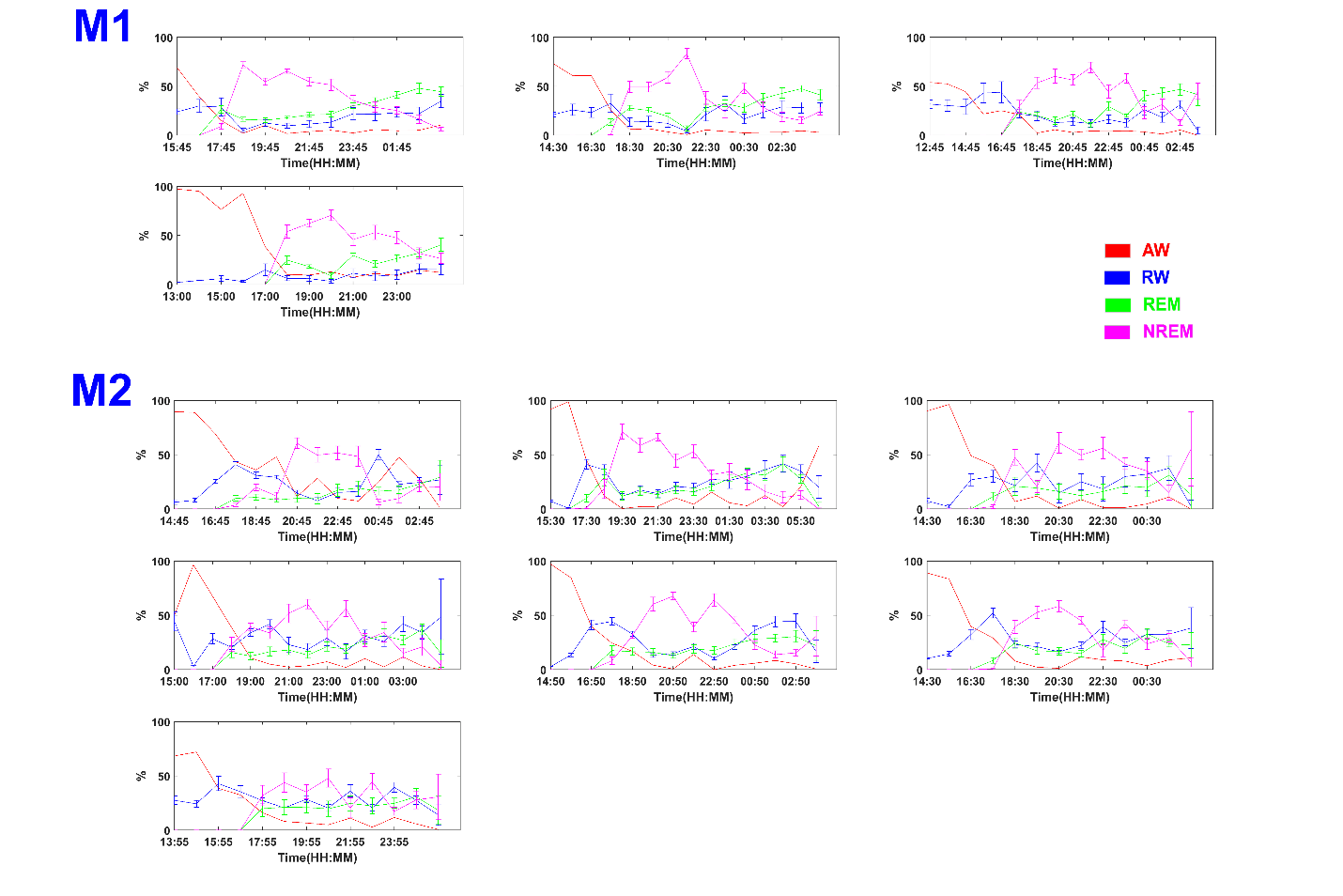

S4: Percentage of time (out of 1 hour) spent in each brain state as a function of time of day. Average (± SD) over all channels. Red: active-wake; blue: resting-wake; green: REM, pink: NREM.

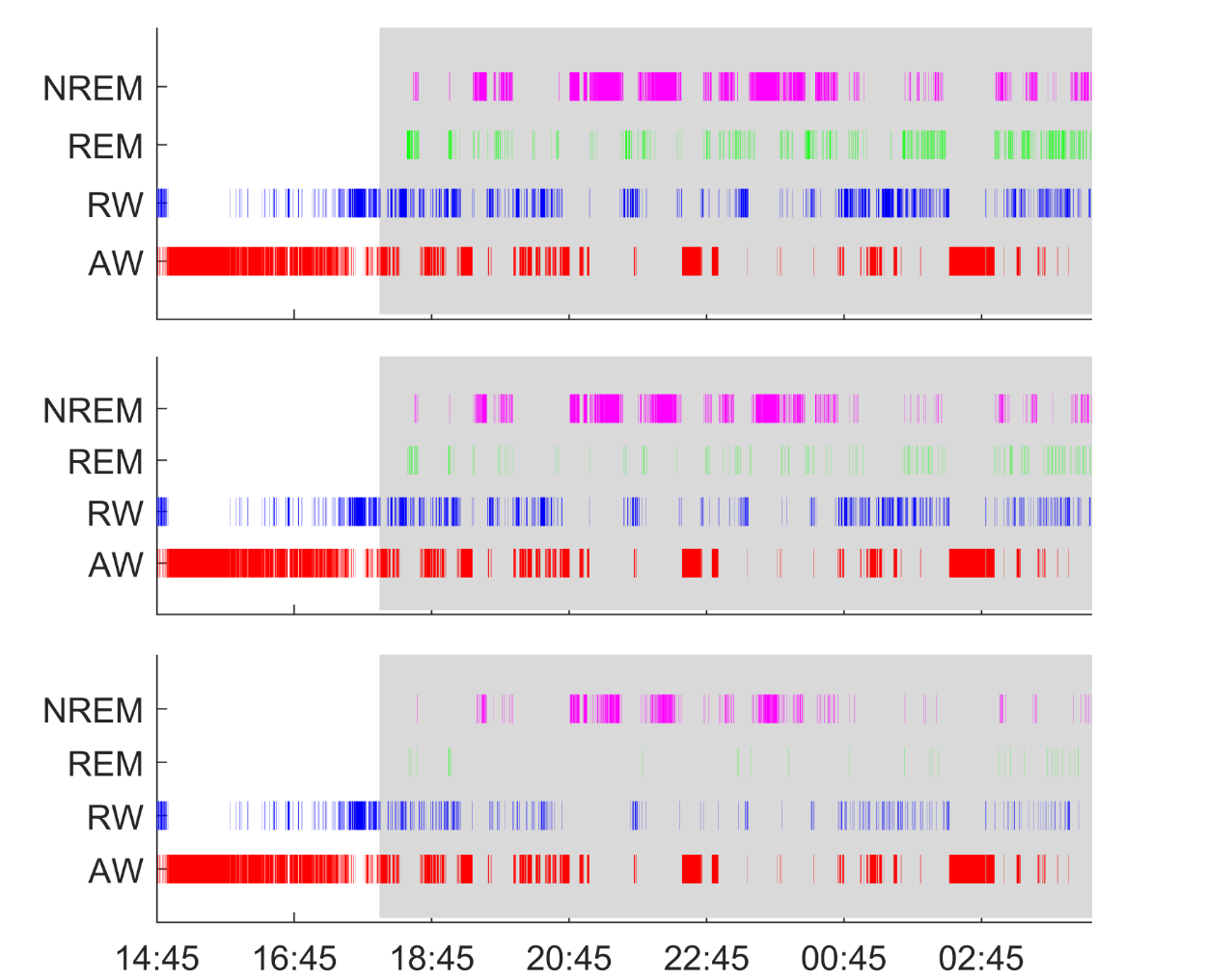

S4B: Brain state classification output for one recording when a minimum of cortical sites are required to report the same brain state in order to assign that state. From top to bottom: less than 5 sites, more than 5 sites, all the sites.

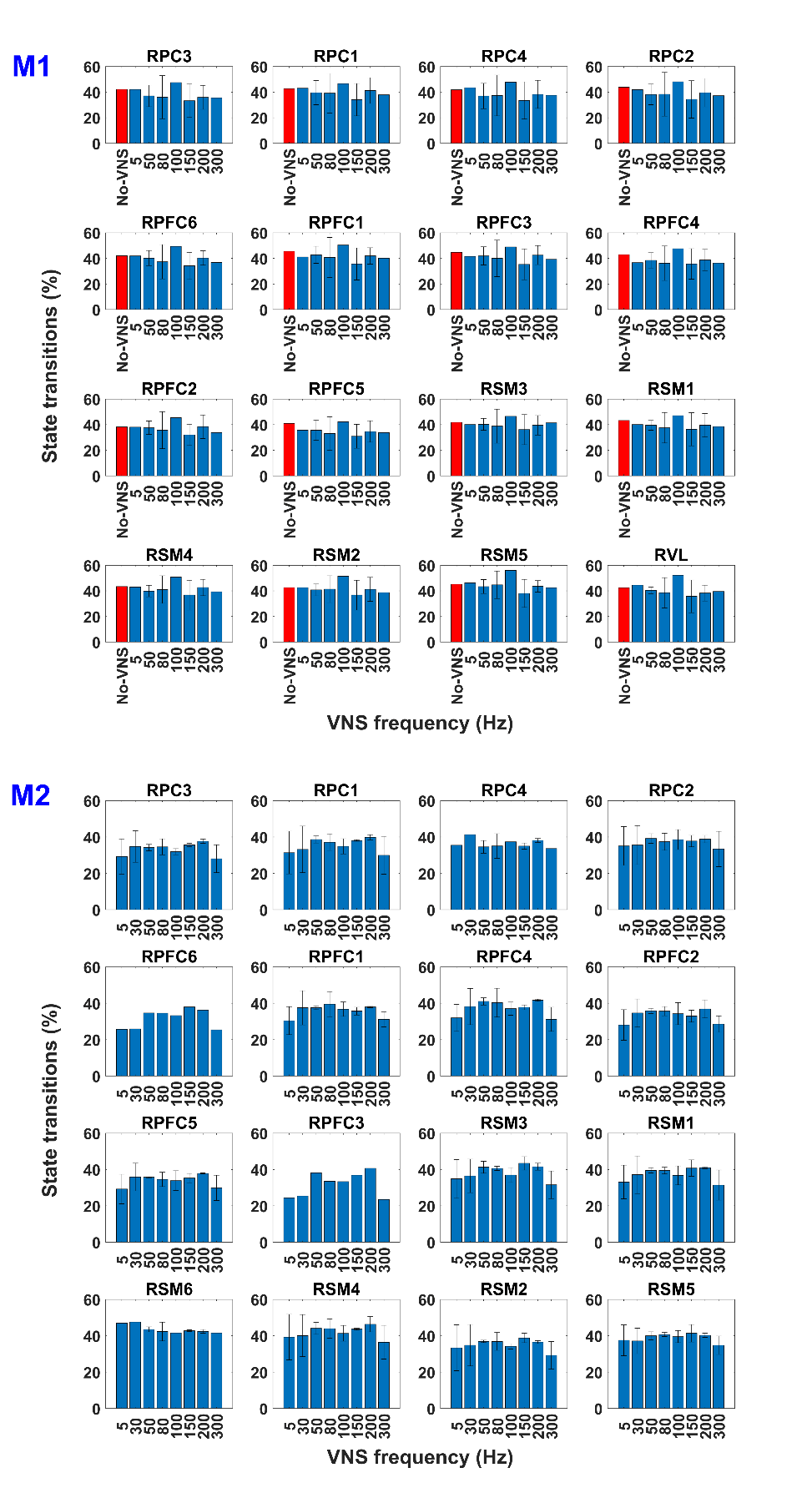

S5: Frequency of brain state transitions before-after VNS across different cortical sites. Each bar represents the % of VNS trains, associated with a brain state transition (brain state after VNS being different than the brain state before VNS). Results are shown for different VNS pulsing frequencies and different cortical sites. The red bar for animal M1 indicates the frequency of state transitions when no stimulation was delivered.

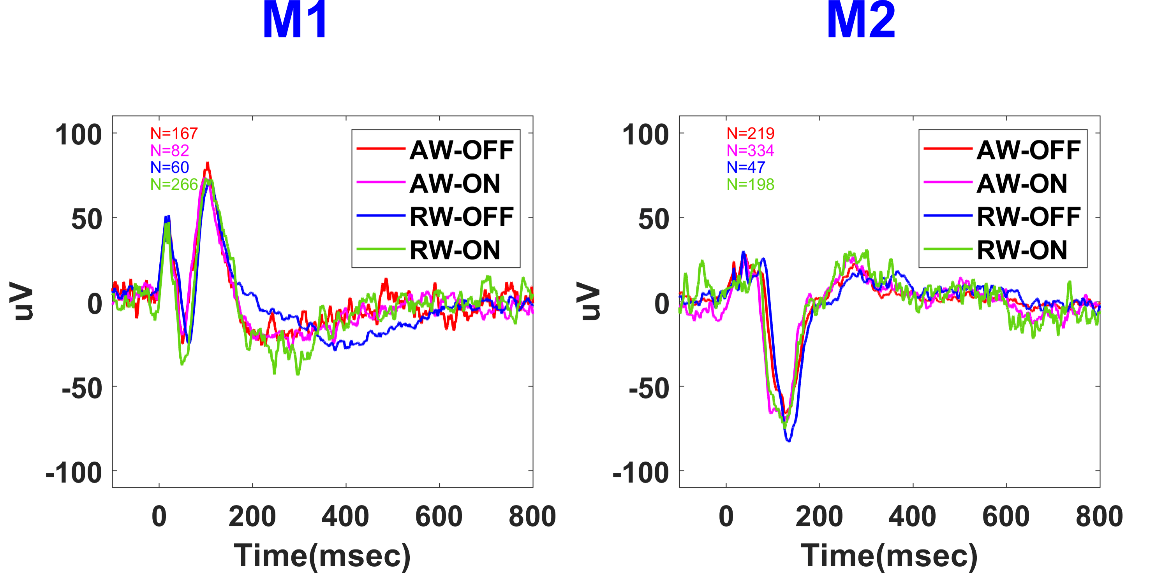

S5B: Comparison of VEPs from one representative recording site classified in the two awake states during light ON and during light OFF for both monkeys. The state of the light (ON or OFF) did not affect the shape nor the time course of the VEPs.

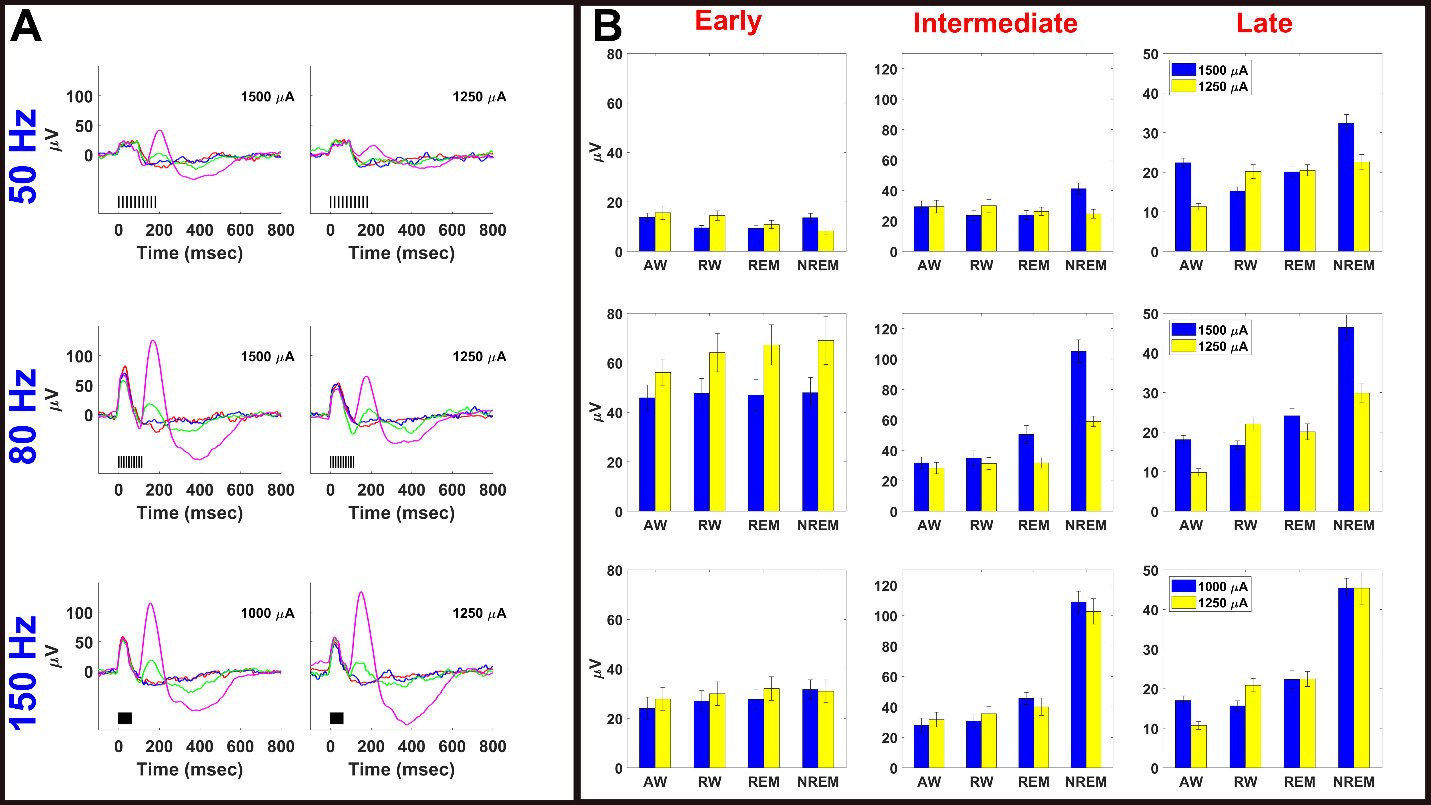

S6: The late components of the VEPs are modulated by brain states independently by the current intensity used to stimulate the vagus nerve for monkey M1. A) Example of classified VEPs from one representative recording site elicited by delivering different current intensities (1250uA, 1000uA and 1500uA) for stimulation frequencies of 50Hz, 80Hz and 150. B) Bar plots showing the averaged absolute value of the maximum deflection of the VEP components over all channels (mean ± SE) evoked by trains of 5 pulses delivered at frequencies of 50Hz, 80Hz and 150Hz as function of brain state (X axis). The two colors represent the two current intensities under comparisons (yellow= 1250 uA, blue= 1000uA or 1500uA). Higher current intensities evoked larger VEPs in all brain states. As shown in Figure 4 the brain-state modulation is larger when higher stimulation frequencies are used.

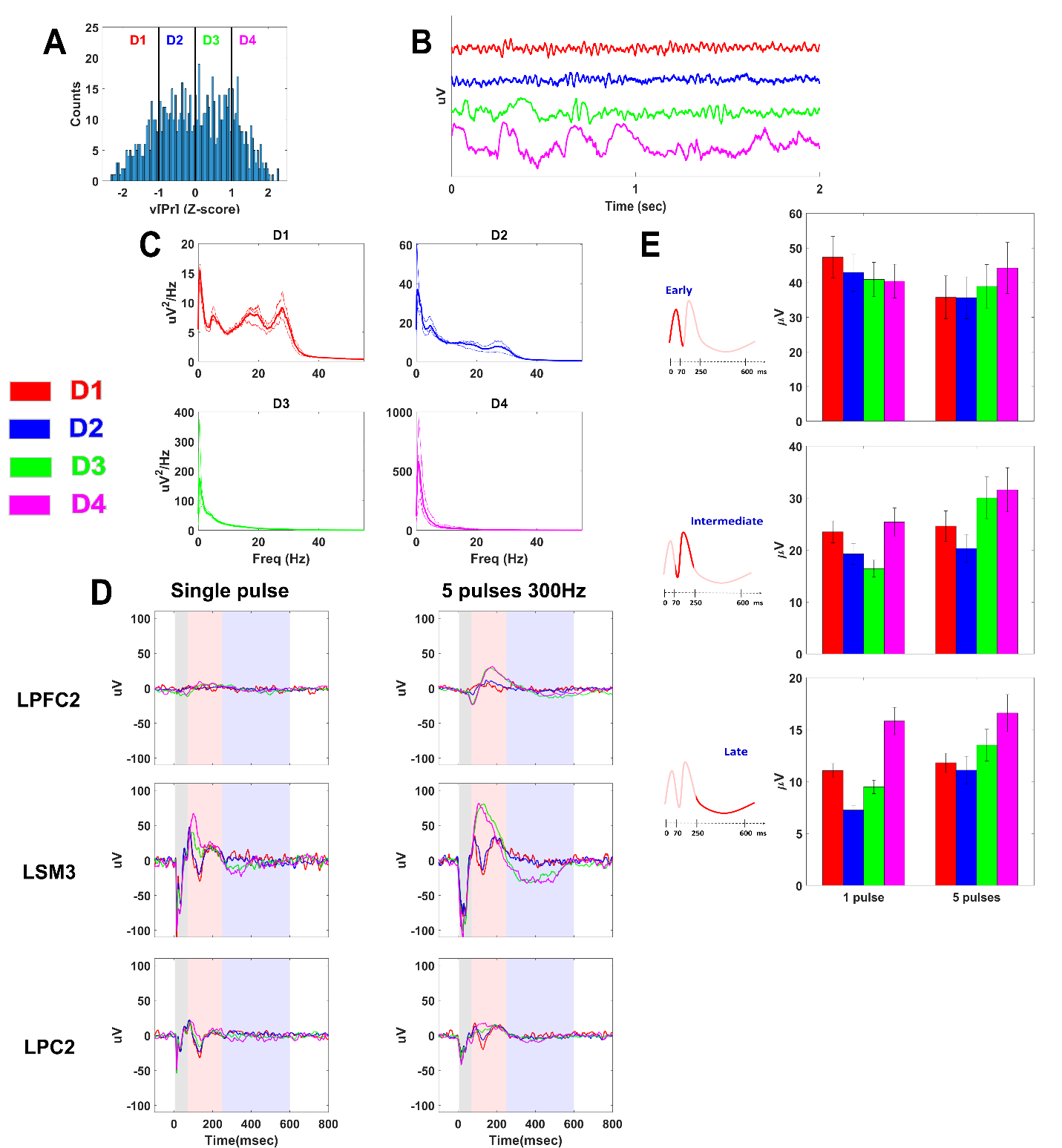

S7: Somatosensory Evoked Potentials (SEPs) elicited by stimulating the median nerve showed a modulation based on delta power contribution in monkey M3. A) Example of Z-scored transformed (see Method) power distributions in delta band generated by all epochs of one recording from one representative cortical site. The epochs were classified based on delta power contribution from D1 (epochs with the lowest delta power) to D4 (epochs with the largest delta power). B) Two seconds of raw signal for each classified delta state of one representative cortical site. C) Power spectrum profile of classified epochs over all cortical channels for each recording (thin traces). Thick traces show the average across the recordings. D) Examples of SEPs elicited during different delta states for three representative sites. SEPs were evoked in one recording by a single pulse at 1500uA (left column) and in a second recording by trains of 5 pulses at 300 Hz (right column). The colored shadow areas represent the time range of each component as defined for the VEPs characterization: early, 5-70ms (gray); intermediate, 70-250ms (orange); late, 250-600ms (light blue). The X axis represents the time after the first pulse in the train. The colored traces represent different brain states: red, D1; blue, D2; green, D3; pink, D4. E) Quantification of the delta state modulation of the three components as averaged of the absolute values of the maximum deflection for each component over all channels (mean ± SE) evoked by a single pulse and trains of 5 pulses at 300Hz.

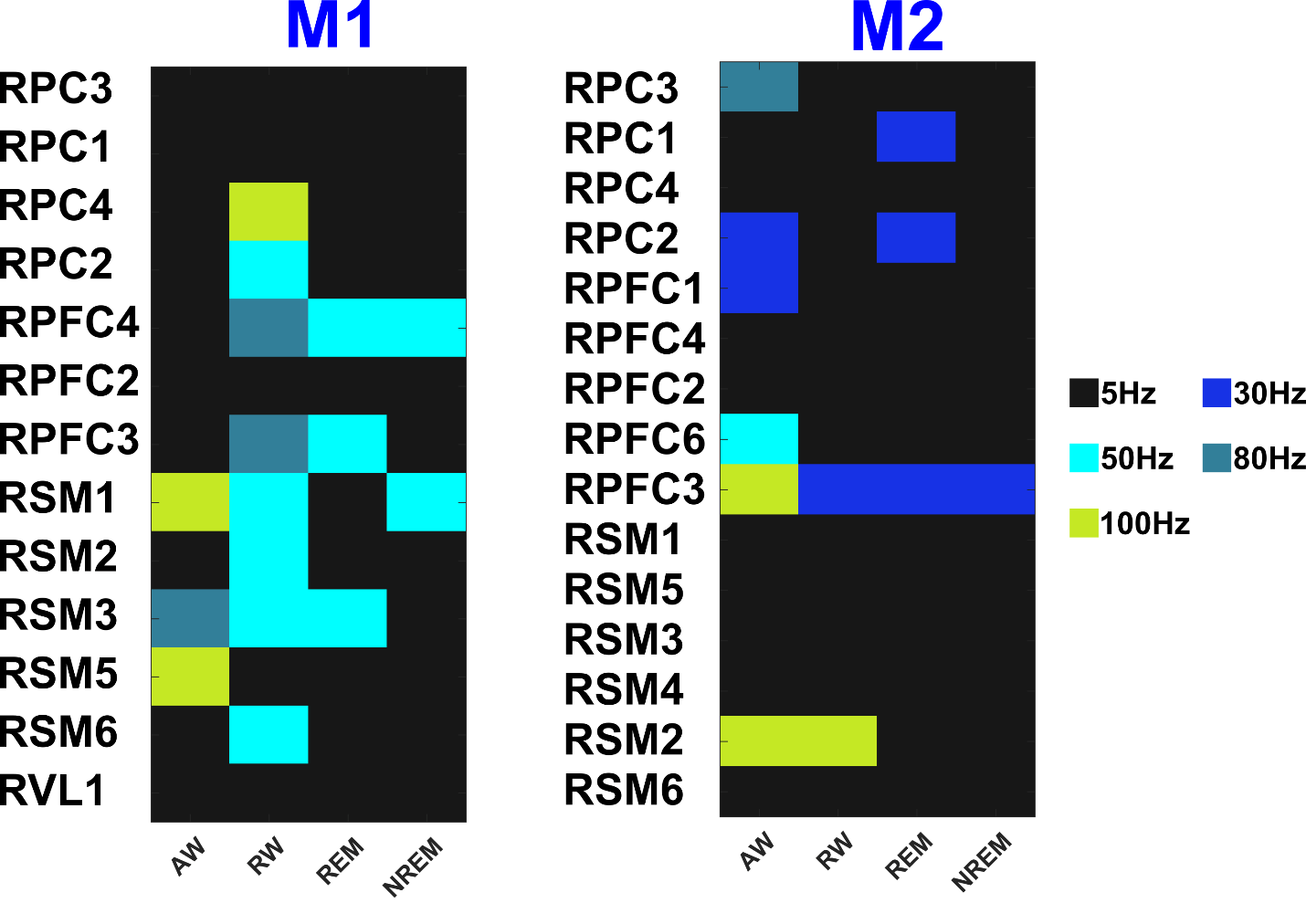

S8: Quantification of the minimum VNS frequency necessary to evoke a significantly large response for each recording site.

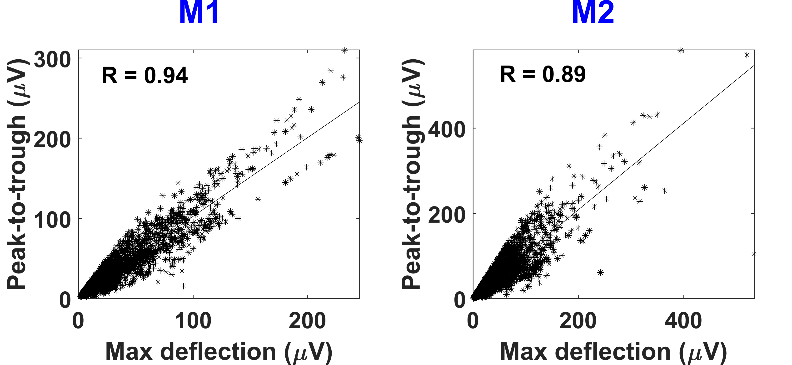

S9: Scatter plot with the least-squares regression line for each animal representing the amplitude of the maximum deflection versus the peak-to-trough magnitude for all three components elicited by all tested VNS protocols. R is the correlation coefficient (p<0.01 for both animals).

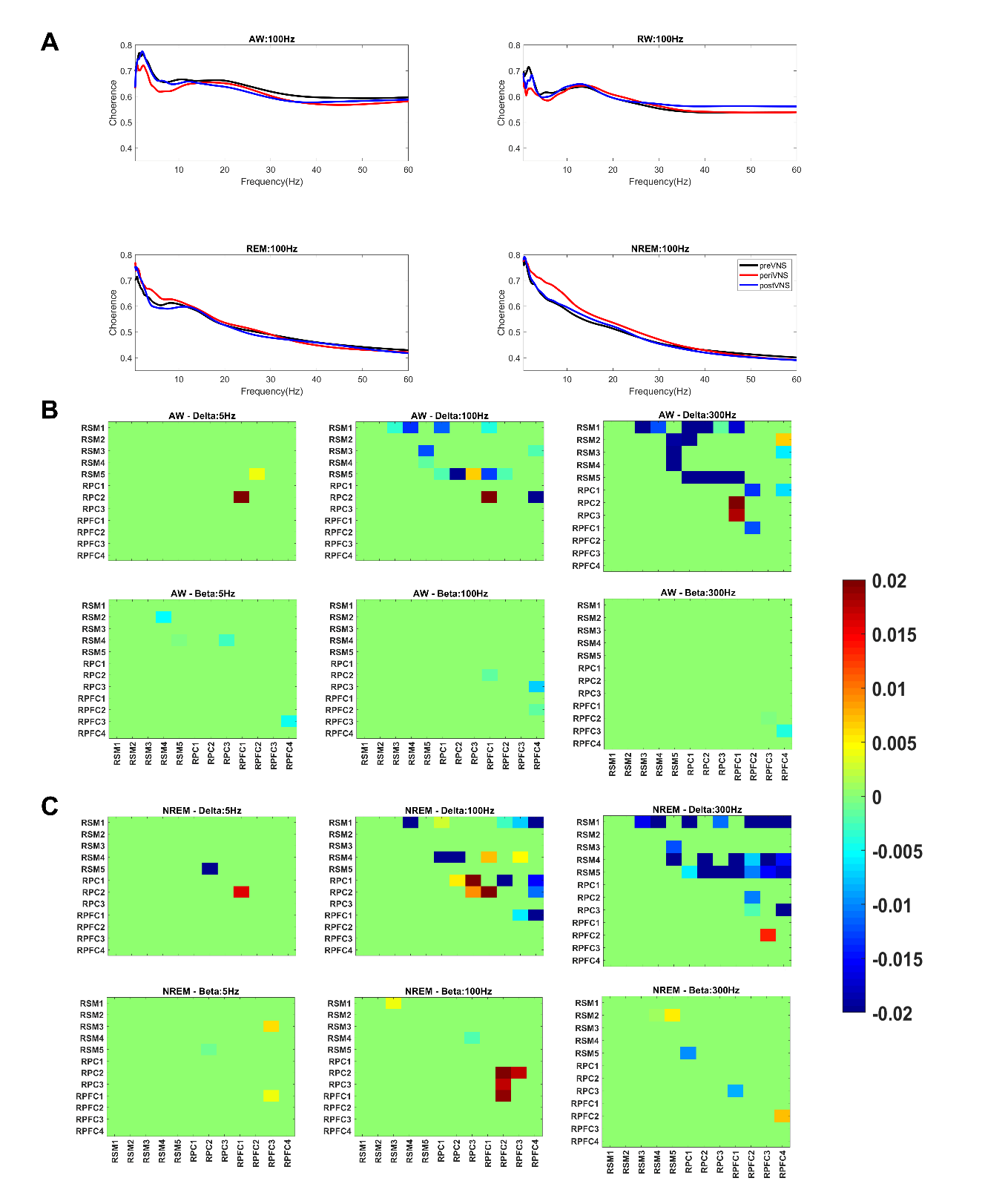

S10: A) Coherence spectrum during active awake(AW), resting awake (RW), REM and NREM sleep for monkey M1 calculated over three different time windows for one pair of channels during 100Hz VNS: 2 seconds before stimulation onset (preVNS, black), 1 second during VNS (periVNS, red) and 2 seconds after VNS in a window between [2-4] sec from stimulation onset (postVNS, blue). B) Color maps showing pairwise significant differences (ttest, p<0.05) in coherence in delta (top) and beta (bottom) range between all channels during awake for 3 different conditions: 5Hz VNS, 100Hz VNS, 300Hz VNS for monkey M2. C) Color maps showing pairwise significant differences (ttest, p<0.05) in coherence in delta (top) and beta (bottom) range between all channels during NREM sleep for 3 different conditions: 5Hz VNS, 100Hz VNS, 300Hz VNS for monkey M2. Refer to Figure 1 for the anatomical location of the electrodes.
